# Redifferentiated cardiomyocytes retain residual dedifferentiation signatures and are protected against ischaemic injury

**DOI:** 10.1101/2022.02.22.481415

**Authors:** Avraham Shakked, Zachary Petrover, Alla Aharonov, Matteo Ghiringhelli, Kfir-Baruch Umansky, Phong Dang Nguyen, David Kain, Jacob Elkahal, Yalin Divinsky, Shoval Miyara, Gilgi Friedlander, Alon Savidor, Lingling Zhang, Dahlia Perez, Nathaniel Kastan, Daria Lendengolts, Yishai Levin, Jeroen Bakkers, Lior Gepstein, Eldad Tzahor

## Abstract

Cardiomyocyte renewal by dedifferentiation and proliferation has fueled the field of regenerative cardiology in recent years, while the reverse process of redifferentiation remains largely unexplored. Redifferentiation is characterised by the restoration of function that is lost during dedifferentiation and is key to the healing process following injury. Previously, we showed that ERBB2-mediated heart regeneration has these two distinct phases: dedifferentiation, followed by redifferentiation. Here, using temporal RNAseq and proteomics, we survey the landscape of the dedifferentiation-redifferentiation process in the adult mouse heart. We find well characterised dedifferentiation pathways, such as reduced oxphos, increased proliferation and increased EMT-like features, largely return to normal, though elements of residual dedifferentiation remain, even after contractile function is restored. These hearts appeared rejuvenated and showed robust resistance to ischaemic injury. We find that redifferentiation is driven by negative feedback signalling, notably through LATS1/2 Hippo pathway activity. Disabling LATS1/2 in dedifferentiated cardiomyocytes augments dedifferentiation *in vitro* and prevents redifferentiation *in vivo*. Taken together, our data reveal the non-trivial nature of redifferentiation, whereby elements of dedifferentiation linger in a surprisingly beneficial manner. This cycle of dedifferentiation-redifferentiation protects against future insult, in what could become a novel prophylactic treatment against ischemic heart disease for at-risk patients.

## Introduction

In recent years, enormous strides have been made towards achieving cardiac regeneration via cardiomyocyte (CM) dedifferentiation and proliferation, however few studies have considered the return journey: how and to what extent do dedifferentiated CMs *redifferentiate*^*1*^? The necessity for redifferentiation has been highlighted by several recent studies whereby irreversible interventions that cause persistent CM dedifferentiation and proliferation proved deleterious in zebrafish^2^ and fatal in mice^3–5^ & pigs^6^. This is consistent with the notion that dedifferentiation endows the adult heart with embryonic characteristics that promote regeneration but hinder mature function such as contractility. Despite this, some studies have actually found that irreversible interventions to induce dedifferentiation are well tolerated by the heart^7^ and improve function after injury^8–15^. We postulate that negative feedback signalling may be the countering force that stabilises or improves heart function in response to mitogenic signalling, and therefore important for redifferentiation. However, its significance has never been investigated, partially hampered by the lack of a suitable model.

To address this, we took advantage of a previously established transgenic mouse model that can transiently express an activated form of the tyrosine kinase receptor ERBB2 (caERBB2)^3^. Such mice show a distinct dedifferentiation phase when caERBB2 is upregulated, characterised by the decline in functional parameters, and a clear redifferentiation phase after caERBB2 expression is halted, characterised by functional recovery^3^. This allowed us to investigate not only the mechanism behind redifferentiation, but to shed light on the extent to which certain biological processes do or do not return to normal after dedifferentiation.

Here, we show that the cycle of ERBB2-induced dedifferentiation and redifferentiation occur independently of injury and that despite functional recovery during redifferentiation, hundreds of genes and proteins remain or become differentially expressed one month after caERBB2 shut-off. These proteins reveal a signature of residual dedifferentiation, whereby certain pathways activated or inactivated during dedifferentiation mostly, but do not completely return to normal. This has the surprising effect of conferring robust cardioprotection to the heart against future ischemic injury, which has the potential to act as a prophylaxis for patients at high risk for MI. Mechanistically, redifferentiation in mice is driven by the Hippo pathway kinases LATS1/2, which when knocked-out in the background of caERBB2 overexpression leads to failed redifferentiation and sustained dedifferentiation. In adult zebrafish, which regenerate naturally, the negative feedback regulators *Dusp6*^*16*^ and *Spry4*^*17*^ are enriched in cycling border zone CMs, suggesting a conserved mechanism for redifferentiation.

The current lack of knowledge around CM redifferentiation is a barrier to clinical translation of regenerative therapies that promote CM dedifferentiation. This study thoroughly characterises the process of redifferentiation and shows that despite the remarkable capacity the heart has to redifferentiate, the process is not trivial, and that dedifferentiation can have lasting effects. In the most extreme case, we found that Connexin43, which is downregulated during dedifferentiation, does not recover for at least one year. A more comprehensive understanding of redifferentiation will be critical in realising the potential for cardiac regeneration.

## Results

### Functional recovery after transient ERBB2 signalling occurs despite incomplete reversal and development of differentially expressed genes and proteins

In order to determine the requirement of transient mitogenic signalling for successful CM redifferentiation during cardiac regeneration, we employed the use of a previously established system to transiently or persistently express caERBB2 in CMs^3,18^. In this paradigm, caERBB2 is activated transiently by temporarily removing Doxycycline (Dox) food, or persistently by permanently removing Dox food (Fig. 1a). Echocardiography of LAD-ligation injured mice revealed that persistent overexpression (pOE) led to a decline in relative stroke volume (rSV), similar to that of a WT mouse, whilst transient OE (tOE) allowed for functional recovery (Fig. 1c). Although the ERBB2-induced hypertrophy was the main driver behind the low rSV, as evidenced by increased wall thickness (Extended Data Fig.1a-b) and decreased LV volume (Fig. 1d), CMs remained dedifferentiated with compromised function even after the hypertrophy subsided, 1 week post caERBB2 shut-off (4WPMI). From this time point until 7WPMI, functional improvement occurred without changes to ventricular dimensions in tOE hearts (Fig. 1c-d (blue shaded area), Extended Data Fig.1a-d). We also found that sham injured hearts showed a very similar functional trajectory, demonstrating the injury-independent nature of the dedifferentiation-redifferentiation cycle (Fig. 1e-f, Extended Data Fig.1e-h). Hence, persistent CM dedifferentiation prevents functional recovery.

**Fig. 1.**
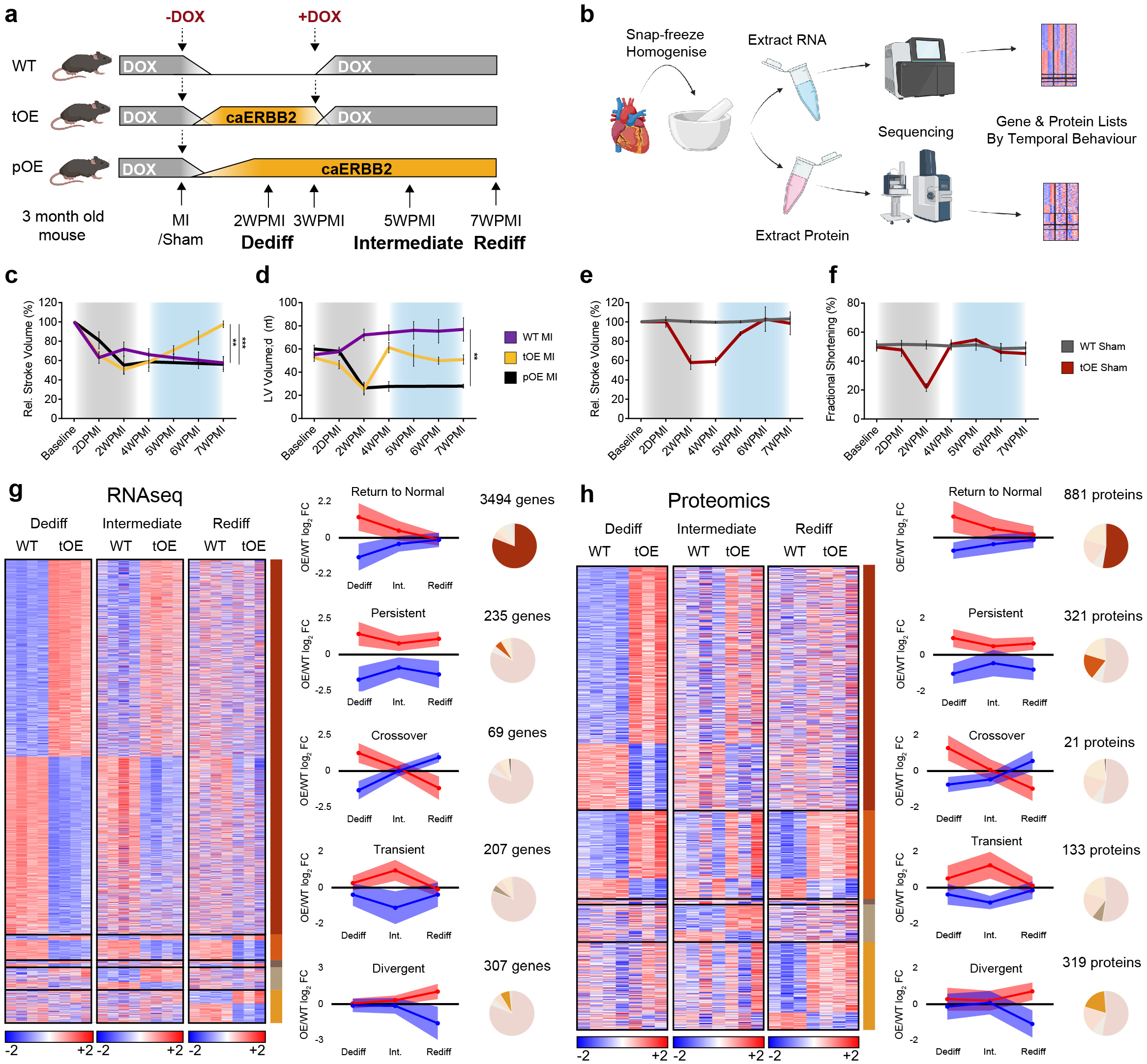
Functional recovery after transient ERBB2 signalling occurs despite incomplete reversal and development of differentially expressed genes and proteins. **a**, Schematic of inducible caERBB2 expression system in adult mouse cardiomyocytes (CMs). -DOX and +DOX represent the respective removal and re-introduction of Doxycycline to the mouse’s diet to temporarily induce caERBB2 expression. **b**, Cartoon schematic of workflow to extract RNA and protein from whole hearts before sequencing. **c-f**, Relative stroke volume **(c**,**e)**, and left ventricular diastolic volume (LV Volume;d) **(d**,**f)**, of hearts from WT Ml, tOE Ml, pOE Ml, WT Sham and tOE Sham mice, measured by echocardiography. Gray shaded area represents the time period of caERBB2 activation (dedifferentiation). Blue shaded area represents the redifferentiation phase. Overall, *n* = 3 - 8 per group. Data are represented as mean± SEM. Statistical significance was calculated by one-way ANOVA with Sidak’s multiple comparison test at the 7WPMI timepoint. *\*p* 0.05, *\*\*p* < 0.01, *\*\*\*p* < 0.001, *\*\*\*\*p* < 0.0001. D/WPMI = days/weeks post myocardial infarction. **g-h**, Heatmaps based on 1092 transformed normalised counts for Sham RNAseq and 10910 transformed intensity values for Sham proteomics. Rows represent genes/proteins. Columns represent each biological sample. Colour bars represent z-score for each row within each timepoint. Graphs show the average tOE/WT 1092 fold change for genes **(g)**, and proteins **(h)**, for each temporal group. Data are represented as mean± SD. Within each temporal group, genes/proteins that were downregulated when first differentially expressed (DE) are shown in blue, and those that were upregulated when first DE are shown in red. Pie charts show the relative proportion of genes/proteins in each temporal group (out of the total number of DE genes/proteins).

Given the requirement of transient dedifferentiation in cardiac regeneration, we sought to survey the transcriptional and proteomic landscape over the course of redifferentiation. We isolated hearts from 3 key time points: peak of ERBB2 activity termed ‘Dediff’, 2-weeks after ERBB2 signalling cessation termed ‘Intermediate’ (Int.), and a point in time where regeneration was estimated to have completed, termed ‘Rediff’ (Fig. 1a). These time points corresponded to a return to baseline values of heart function (Fig. 1c-f) and ERBB2 levels (Extended Data Fig.1i-k). We performed bulk RNAseq and proteomics on sham and injured, WT and tOE hearts, from these three timepoints (Fig. 1b), and procured a list of genes or proteins that were differentially expressed at at least one time point. We grouped the genes and proteins according to 5 different temporal patterns; Return to normal, Persistent, Cross-over, Transient and Divergent (Fig. 1g-h). Over the time period of functional restoration, (for both Sham and MI hearts), the majority of these genes and proteins showed a ‘Return to Normal’ pattern, however a significant proportion remained, or actually became differentially expressed i.e. ‘Persistent’ or ‘Divergent’ respectively (Fig. 1g-h, Extended Data Fig. 1l-m). This is reflected in the PCA plots and corresponding dendrograms (Extended Data Fig. 1n-q). Hence, functional recovery after transient ERBB2 signalling occurs despite incomplete reversal and development of differentially expressed genes and proteins.

### Dedifferentiation phenotypes are largely reversed in functionally redifferentiated hearts

We next sought to understand whether the differentially expressed genes and proteins at the ‘Rediff’ timepoint were clustered into pathways that could reveal previously unknown functional differences between tOE and WT hearts. To do so, we performed Ingenuity Canonical Pathways analysis on the sham RNAseq data. Given the similarity of the sham and MI-injured groups, both in their functional, and gene/protein expression response to transient ERBB2 signalling, we focused only on sham or uninjured animals, unless otherwise stated. By plotting the enrichment significance (tOE/WT) against the z-score for GO terms at all 3 timepoints (Dediff, Int. and Rediff), we were able to visualise canonical pathway behaviour over time. We found that well characterised dedifferentiation signatures, such as increased proliferation^3,4^, increased EMT-like features^9,18^ and decreased oxphos metabolism^19,20^ all returned to normal (Fig. 2a). We validated this by RT-qPCR, also assaying for the lactate transporter *Slc16a3* indicating a complementary shift to glycolysis (Extended Data Fig. 2a) and noted that when ERBB2 signalling was persistent, these genes remained differentially expressed (Extended Data Fig. 2b). We also found that by western blot, many proteins involved in these pathways showed similar expression trajectories (Fig. 2b, Extended Data Fig. 2c). Since data from bulk-sequencing methods can be influenced by various cell types in the heart, we validated these signatures were relevant for CMs by staining *in vivo* heart sections (Fig. 2c). To determine whether the Ingenuity pathway analysis prediction of oxphos metabolism inhibition at the Dediff time point (Fig. 2a) translated into a measurable change in metabolism, we cultured CMs from P7 WT and OE pups (as an *in vitro* proxy for the Dediff timepoint). Accordingly, we found that the OE CMs had diminished maximal oxygen consumption and increased ECAR (proxy for glycolysis) (Extended Data Fig. 2d). Furthermore, we also found that hypercellularity and gross tissue disarray, typical of dedifferentiated hearts^3^, also returns to normal by the Rediff timepoint (Extended Data Fig. 2e).

**Fig. 2.**
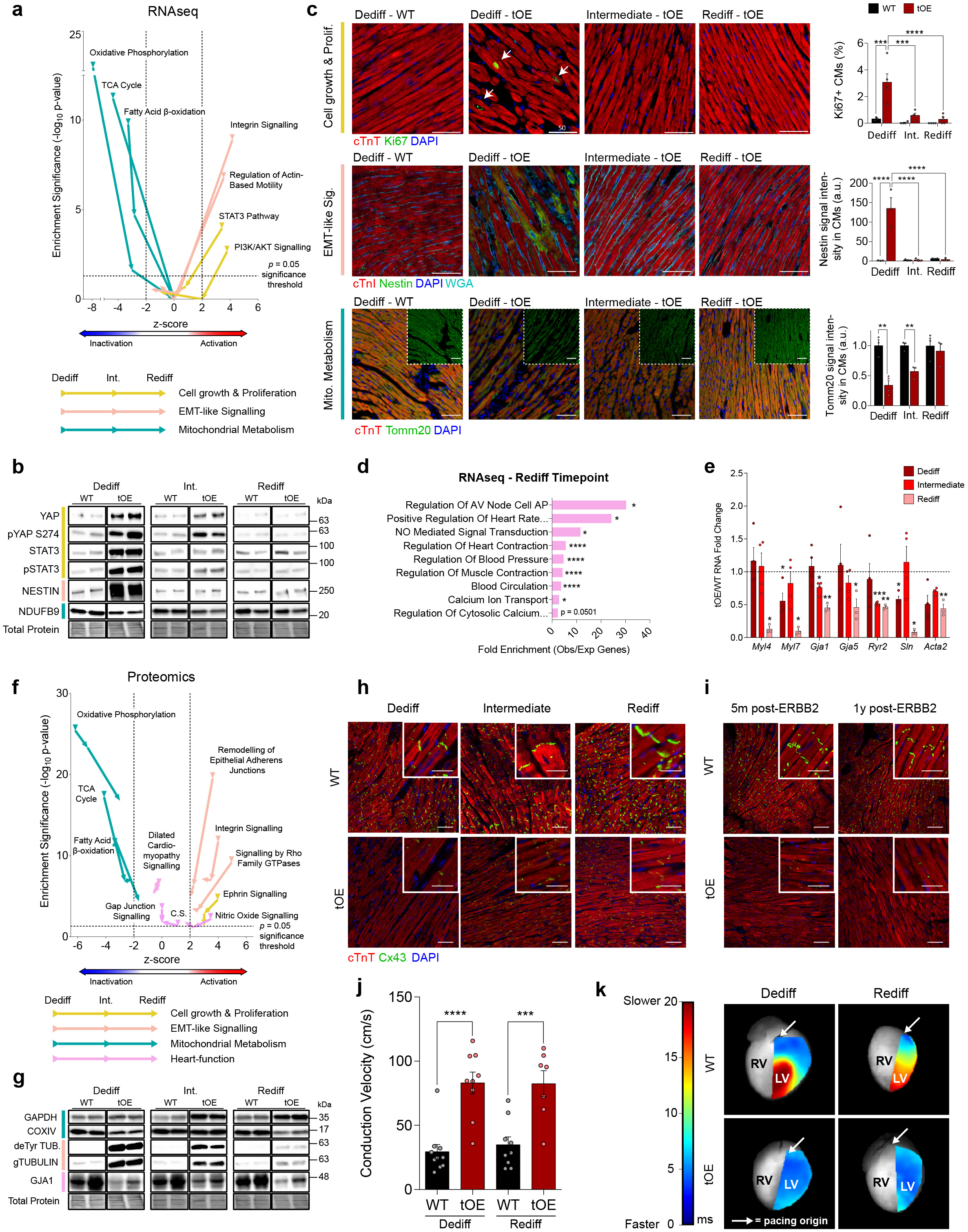
Dedifferentiated phenotypes are largely reversed in functionally redifferentiated hearts. **a** Scatter plot of GO term z-score against enrichment significance (log_10_ p-value) for tOE/WT across all timepoints, based on Ingenuity Canonical Pathway Analysis of RNAseq data using a threshold fold change (FC) ≥ 1.5; adjusted p ≤ 0.05. Arrows on each line indicate the direction of the GO term from Dediff to Int. to Rediff. Z-scores below -2 are predictive of pathway inactivation and above +2 are predictive of pathway activation. Values above the horizontal dashed black line represent statistically significant enrichment. **b** Representative western blot of ‘Return to normal’ proteins for WT and tOE adult heart lysates for each time point. *n* = 4 - 7 per group. **c** Representative immunofluorescence images of WT hearts at Dediff, and tOE hearts at Dediff, Int. and Rediff, with full quantification. Images were acquired in the remote zones of MI injured hearts as a proxy for sham injury. White arrows highlight Ki67+ CMs. *n* = 3 - 5 per group. Scale bars = 50μm for Ki67 and Nestin, 100μm for Tomm20. **d** Panther Over-representation results for tOE/WT differentially expressed (DE) genes at Rediff. **e** RT-qPCR analysis of heart-function related genes that are DE at Rediff. Each value is normalised to the WT value of its corresponding time point. **f** Equivalent scatter plot to (**a**), based on proteomic data, using a threshold FC ≥ 1.1; p ≤ 0.05. C.S. = Calcium Signalling **g** Representative western blot for WT and tOE adult heart lysates at each time point for proteins that were DE at Rediff. **h-i** Immunofluorescence images of Cx43 in adult WT and tOE hearts at all timepoints **(h)** plus additional timepoints of 5 months and 1 year post-ERBB2 shut-off **(i)**. Main scale bar = 100μm, inset 10μm. **j** Conduction velocity measured by high-resolution optical mapping of epicardial action potential across the left ventricle of langendorff perfused hearts. *n* = 7 - 10 per group, paced from the base of the heart every 200ms. **k** Representative heatmaps of conduction velocity. Each pixel is coloured according to the amount of time taken for the action potential (originating at the pacing electrode) to reach it, overlaid onto a greyscale image of the heart. RV = Right Ventricle. LV = Left Ventricle. In all panels numerical data are presented as mean ± SEM; statistical significance was calculated using two-way ANOVA followed by Sidak’s test in **(c)** for Ki67 and Nestin and in **(j)**, two-tailed unpaired Student’s t-test in **(c)** for Tomm20 and **(e)** for WT to tOE comparison within each time point and Fisher’s Exact test with the Benjamini-Hochberg false discovery rate (FDR) correction for multiple testing in **(d)**. \**p* ≤ 0.05, \*\**p* < 0.01, \*\*\**p* < 0.001, \*\*\*\**p* < 0.0001.

Although the above data are consistent with the return to normal contractility during redifferentiation, further analysis reveals a more complex picture, whereby some canonical pathways remain significantly enriched at the Rediff timepoint. For example, Panther analysis of differentially expressed genes at Rediff revealed enrichment of several pathways relating to heart function (Fig. 2d). The gene expression changes driving this result (including *Myl4, Gja1* and *Ryr2*) were validated by RT-qPCR (Fig. 2e). Ingenuity Canonical Pathway analysis of the proteomic data complemented this, with persistent enrichment for the GO terms: Dilated Cardiomyopathy Signalling, Nitric Oxide Signalling, Gap Junction Signalling and Calcium Signalling (Fig. 2f). This analysis also revealed that at the Rediff timepoint, pathways involved in proliferation, EMT-like signalling and mitochondrial metabolism retained a significant activation and inactivation prediction respectively (Fig. 2f). These predictions are driven by a concentration of differentially expressed proteins in these pathways at the Rediff timepoint (Extended Data Fig. 2f-h), even if the expression fold change is less dramatic (COXIV, TUBULIN & deTyrosinated TUBULIN) (Fig. 2g, Extended Data Fig. 2i). The trajectory of these pathways appears to indicate a residual level of dedifferentiation that remains at the Rediff timepoint, however the heart function related pathways (marked in pink) are more static. One such protein driving these predictions was the gap junction protein Cx43, which was persistently downregulated in response to ERBB2 signalling (Fig. 2g, Extended Data Fig. 2i). Given its important role in the heart^21^, we stained WT and tOE sections from all 3 timepoints and found that both its intensity and gap junction localisation were markedly reduced (Fig. 2h). We posited that if given more time to redifferentiate, Cx43 levels might recover. Therefore we stained heart sections that were 5 months and 1 year post ERBB2 shut-off, and remarkably the levels were still drastically lower than the WT, indicating that there are some potentially irreversible consequences of CM dedifferentiation (Fig. 2i).

Although by echocardiography, WT and tOE hearts at the Rediff timepoint are indistinguishable, we hypothesised that conduction velocity, partially mediated by Cx43, would reveal a difference between the two groups. To test this, we performed *ex vivo* high-resolution optical mapping of the action potential across the subepicardium in WT and tOE hearts at the Dediff and Rediff timepoints. We found that tOE hearts accommodated a significantly faster conduction velocity, revealing a previously unknown functional consequence of the Rediff state (Fig. 2j-k), that could not be detected by echocardiography. Although a reduction in Cx43 is commonly associated with a reduction in conduction velocity, this typically only occurs after a more than 90% reduction in total levels^22^ (beyond what we measured) but also can be influenced by various other factors^23^. Taken together, the molecular signatures of dedifferentiation are largely but not completely reversed during redifferentiation.

### Dedifferentiation-Redifferentiation cycle confers robust cardioprotection against ischaemic injury

Given that redifferentiated hearts have normal echocardiographic function with remnant molecular features of embryonic or neonatal hearts (e.g. proliferation, metabolism and EMT-like features) we sought to test whether a cycle of dedifferentiation-redifferentiation (DR): 3 weeks of caERBB2 ON, followed by 4 weeks of caERBB2 OFF, rejuvenates the heart and consequently confers greater resilience to injury. We therefore performed LAD ligation (MI) on WT and tOE-DR adult mice i.e. those that had undergone a cycle of DR, and recorded their heart function for 4 weeks (Fig. 3a). Whilst WT heart function parameters declined after injury, with a wide distribution of values, tOE-DR mice retained near-baseline values, with a tighter distribution (Fig. 3b-d). This effect was seen as early as the 2-day post MI time point, and persisted for a month, indicating a robust cardioprotective effect of a DR cycle before injury. This effect was predominantly driven by a maintenance of anterior wall thickness (Fig. 3e) and contractility (Fig. 3f), as opposed to a change in posterior wall thickness or ventricular dimensions, which can influence functional echocardiographic parameters (Extended Data Fig. 3a-b). These functional data were complemented by scar type and scar area analysis. tOE-DR hearts had a higher proportion of sections per slide with no scar and a lower proportion with a transmural scar (which in humans correlates with an increased risk of acute mortality following MI^24^) (Fig. 3g,i). Similarly, scar area as a proportion of the left ventricle was lower amongst the tOE-DR group (Fig. 3h-i).

**Fig. 3.**
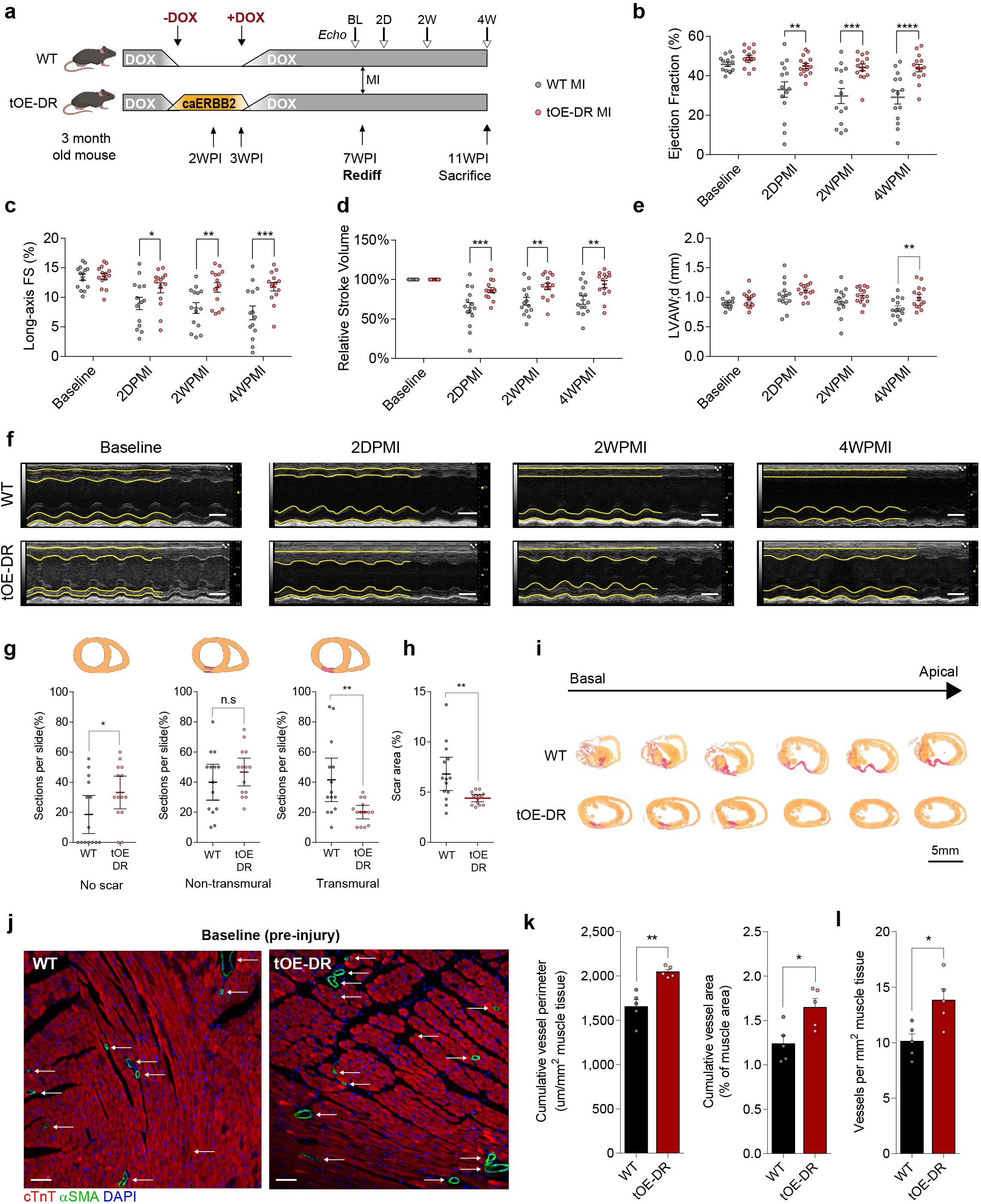
Dedifferentiation-Redifferentiation cycle confers robust cardioprotection after ischaemic injury. **a** Schematic of experimental layout. MIs were performed on WT and tOE-DR mice at the Rediff timepoint. PI = Post Induction (of caERBB2). BL = Baseline, 2D = 2 days, 2W = 2 weeks, 4W = 4 weeks. White arrowheads represent time points for echocardiography. **b-e** Ejection Fraction **(b)**, long-axis Fractional Shortening **(c)**, Relative stroke volume (rSV) **(d)**, Left Ventricular Anterior Diastolic Wall thickness (LVAW;d) of WT and tOE-DR mice, measured by echocardiography. *n* = 14 for each group. **f** Representative M-mode images at the indicated time points. Yellow lines trace wall contractility. Top wall = anterior, bottom wall = posterior. Scale bars = 2mm. **g-i** scar type quantification (**g**), scar area quantification (**h**) and sequential Sirius Red-stained sections from representative WT and tOE-DR hearts 4WPMI (**i**). *n* = 14 for each group. **j-l** Representative ?SMA stained sections highlighting blood vessel density and size distribution (**j**) scale bars = 100μm, cumulative vessel perimeter (left panel) and area (right panel) (**k**) and vessel density (**l**) for WT (27.5mm^2^ tissue quantified across *n* = 5) and tOE-DR (29.3mm^2^ tissue quantified across *n* = 5). In all panels numerical data are presented as mean ± SEM; statistical significance was calculated using two-tailed unpaired Student’s t-test in (**b-e**), (**g**) for non-transmural and transmural scar, (**h**) and (**k-l**) between the in-time point WT and tOE values, and a one-tailed Mann-Whitney test in (**g**) for No scar. \**p* ≤ 0.05, \*\**p* < 0.01, \*\*\**p* < 0.001, \*\*\*\**p* < 0.0001.

Given the previously reported effects of ERBB2 signalling on angiogenesis during injury, we wondered whether tOE-DR hearts had enhanced vascularisation before injury. To that end we stained WT and tOE-DR uninjured heart sections for ?SMA and found increased cumulative blood vessel perimeter and area (Fig. 3j-k). This was driven by an increase in blood vessel density (Fig. 3l), whilst the distribution of vessel size remained unchanged between both groups (Extended Data Fig. 3c-d). This may be one possible mechanism of the observed cardioprotection, which is covered further in the discussion. Taken together, these results indicate that the differences between tOE-DR and WT hearts at the Rediff timepoint are sufficient to confer a powerful cardioprotection against ischemic injury.

### ERBB2 signalling promotes a multi-faceted negative feedback response

Although we demonstrated how tOE-DR and WT hearts behave very differently following injury, the fact that they behave so similarly under basal conditions is a testament to the remarkable journey they undergo from a state of dedifferentiation to redifferentiation. We next asked what could drive such a dramatic phenotypic change over the course of redifferentiation. We previously showed that CM-restricted ERBB2 activation could drive Yap-nuclear entry, despite Hippo pathway upregulation^18^. This led us to hypothesise that the Hippo pathway is being activated as part of a negative feedback loop, and drives the redifferentiation process. Accordingly, we found that the Hippo pathway was activated (upregulation of MST1, LATS1, pLATS1, MOB1, pMOB1, pYAP 112) during the Dediff time point and gradually returned to normal by the Rediff timepoint (Fig. 4a-b, Extended Data Fig. 4a), reflecting an alleviation in negative feedback. Given the previously reported role of ERK and AKT in ERBB2-mediated CM dedifferentiation^3,18^, we broadened our investigation to their negative feedback regulators, amongst others. Analysis of our RNAseq data revealed an activation and ‘Return to Normal’ behaviour in many of these negative feedback regulators, which have reported tumour suppressor activity, including *Ptprv*^*25*^, *Ptprf*^*26*^, *Ptpn9*^*27*^, *Ptpn23*^*28*^, *Phlda1*^*29*^, *Phlda3*^*30*^, *Dusp5*^*31*^, *Dusp6*^*32,33*^, *Spry4*^*17*^, *Numbl*^*34*^, and *Nab2*^*35*^ (Fig. 4c-d, Extended Data Fig. 4b). In order to determine whether or not this finding was unique to our transgenic system, or occurred naturally, we analysed data from previously published data sets^36,37^. We found that just as the expression of negative feedback regulators subsides when caERBB2 signalling is halted in our transgenic system, so too does their expression subside in WT mouse hearts from P0 to adulthood, as ERBB2 expression declines (Extended Data Fig. 4c)^36^. A similar pattern is also seen when only the myocyte fraction of the heart is assayed (Extended Data Fig. 4d)^37^.

**Fig. 4.**
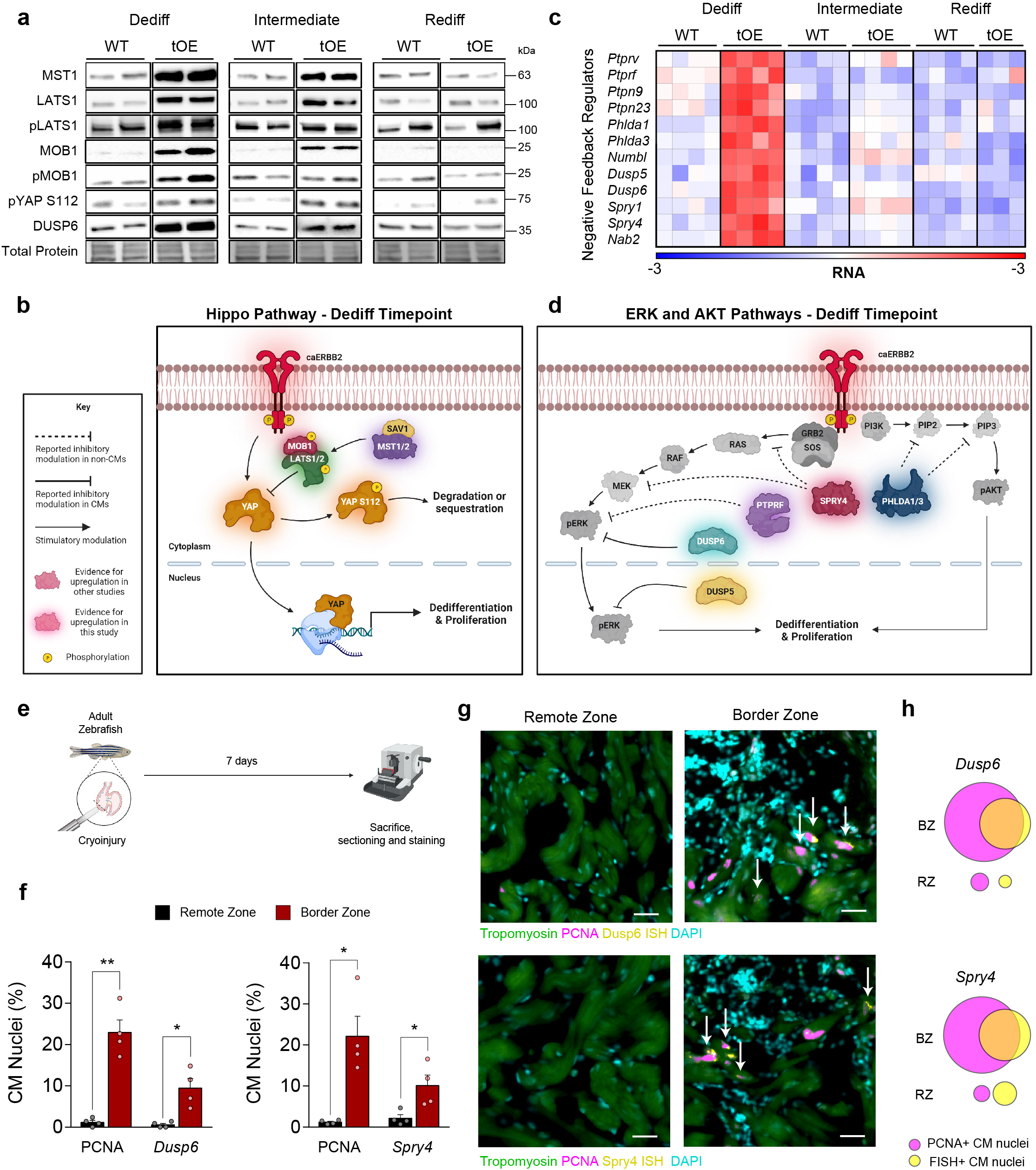
ERBB2 signalling promotes a multi-faceted negative feedback response. **a** Representative western blot of whole-heart lysates for Hippo pathway components and Dusp6 for WT and tOE adult heart lysates for each time point. b Schematic of the Hippo pathway, incorporating western blot data from the Dediff timepoint. **c** Heatmap of negative feedback (NF) regulators, based on log, transformed normalised counts from RNAseq data. Rows represent genes. Columns represent each biological sample. Colour bars represent z-score for each row across all timepoints. **d** Schematic of ERK and AKT pathways, incorporating RNAseq and RT-qPCR data. **e** Experimental plan for assaying NF regulation of zebrafish border zone CMs. **f** Quantification of combined FISH and immunofluorescence staining for PCNA and either *Dusp6* (left panel) or *Spry4* (right panel) positive CM nuclei in remote and border zones of 7DPI adult zebrafish hearts *(n* = 4). **g** Representative images of remote and border zones for *Dusp6* (upper) and *Spry4* (lower) stained sections. White arrows indicate double-positive CM nuclei. Scale bars represent 20µm. **h** To-scale Venn diagram for *Dusp6* (upper) and *Spry4* (lower) positive nuclei overlapping with PCNA positive nuclei between the remote zone (RZ) and border zone (BZ). In all graph panels numerical data are presented as mean± SEM; statistical significance was calculated using a two-tailed paired Student’s t-test in **(f)** between the corresponding remote and border zone values. *\*p* 0.05, *\*\*p* < 0.01, *\*\*\*p* < 0.001, *\*\*\*\*p* < 0.0001

Finally, we wondered whether this phenomenon was conserved in adult zebrafish, which readily regenerate their hearts after injury^38^. To answer this, we performed cryoinjury in adult zebrafish hearts and collected them for sectioning at 7 days post injury (DPI), when CM proliferation peaks (Fig. 4e). A low-throughput screen using *in situ* hybridisation revealed that the Hippo pathway genes *Lats1, Lats2*, and *Sav1*, were not upregulated at the border zone (BZ), but the ERK1/2 negative feedback regulators *Dusp6* and *Spry4* were (Extended Data Fig. 4e-f), suggesting that the zebrafish may rely on different negative feedback mechanisms, or that the Hippo pathway is activated post-translationally. In order to see whether or not this elevated expression was enriched in cycling CMs, we performed fluorescent *in situ* hybridisation (FISH) for both *Dusp6* and *Spry4* along with immunofluorescence for PCNA. As expected, we found that both genes and PCNA were enriched in the border zone (Fig. 4f). Crucially however, we found within the border zone that 87% of *Dusp6* positive CM nuclei and 77% of *Spry4* positive CM nuclei also expressed PCNA (Fig. 4g-h), indicating that their expression is linked to an increase in cycling activity. Hence, mitogenic signalling in mouse and zebrafish CMs promotes a potent negative feedback response.

### Disabling LATS1/2 negative feedback signalling augments dedifferentiation and prevents redifferentiation

In order to interrogate the biological function of negative feedback signalling in the context of redifferentiation we treated WT and ERBB2-OE P7 cardiac cultures with the novel LATS1/2 inhibitor, TRULI^39^ (Fig. 5a). To capture true proliferation events (i.e. with successful cytokinesis) we performed time-lapse microscopy of tdTomato labelled CMs^18^ and validated with immunofluorescence. We found that whilst TRULI had a negligible effect on the proliferation of WT CMs, it caused a dramatic increase in the already proliferative OE CMs (Fig. 5b). This pattern was consistent with staining for AurkB and Ki67 (Fig. 5c, Extended Data Fig. 5a), suggesting that the Hippo pathway actively suppresses ERBB2 driven cell cycle activity, and that Hippo inhibition leads to augmented dedifferentiation.

**Fig. 5.**
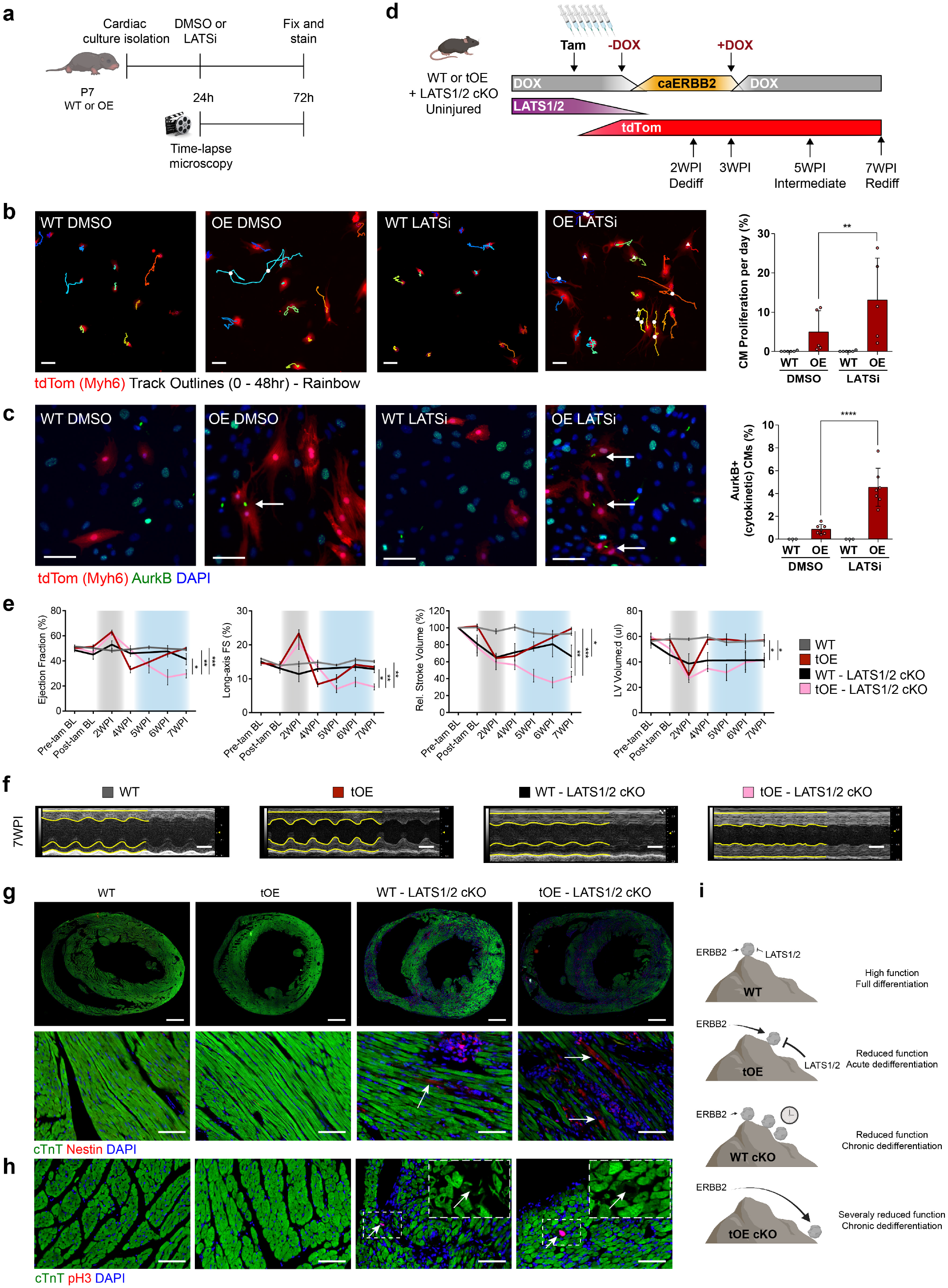
Disabling LATS1/2 negative feedback signalling augments dedifferentiation and prevents redifferentiation. **a** Schematic of experimental layout. **b** Representative still images from first frame of 48hr time-lapse movies of LATSi or DMSO treated WT and OE P7 CMs, with full quantification (*n* = 5 - 6 for each group). White triangles indicate mitosis without cytokinesis and white circles indicate mitosis with cytokinesis. Coloured tracks outline CM displacement. Scale bars = 100 µm. **c** Representative immunofluorescence images for AurkB of groups stated in (**b**) with full quantification. All groups had minimum *n* = 3. Scale bars = 50μm. **d** Schematic of inducible caERBB2 expression system in adult mice with Tamoxifen inducible LATS1/2 cKO and tdTomato expression. **e** Panels from left to right show ejection fraction, Long-axis fractional shortening (FS), Relative stroke volume and left ventricular diastolic volume (LV Volume;d) of WT, tOE, WT LATS1/2 cKO and OE LATS1/2 cKO uninjured mice, measured by echocardiography. Gray shaded area represents the time period of caERBB2 activation. Blue shaded area represents the redifferentiation phase. *n* = 3 - 8 for each group. **f** Representative M-mode images of the left ventricle in diastole and systole for WT, tOE, WT cKO and OE cKO uninjured mice at Rediff. Yellow lines trace wall contractility. Scale bars = 2mm. **g-h** Representative immunofluorescence images of WT, tOE, WT cKO and OE cKO uninjured hearts at Rediff for Nestin (**g**) and pH3 (**h**). For (**g**), upper panel scale bars = 1mm. For lower panel and (**h**), scale bars = 50μm. Arrows highlight CMs positive for either Nestin or pH3. **i** Metaphorical model for the role of negative feedback signalling in redifferentiation. In all panels numerical data are presented as mean ± SEM; statistical significance was calculated using a paired two-way ANOVA followed by Sidak’s test in (**b-c**) and a one-way ANOVA followed by Tukey’s test in the 3 left-most panels of (**e**) and Bonferroni’s test for the following hypotheses: tOE vs. WT cKO and tOE vs. OE cKO in the right-most panel of (**e**). \**p* ≤ 0.05, \*\**p* < 0.01, \*\*\**p* < 0.001, \*\*\*\**p* < 0.0001.

We next asked whether disabling LATS1/2 played a role in suppressing ERBB2-mediated dedifferentiation *in vivo* and by extension would prevent redifferentiation. To that end we generated a mouse model that allows for CM-restricted LATS1/2 deletion^8^ (with tdTomato lineage tracing) using a *Myh6* driven tamoxifen-inducible Cre, in the background of transient ERBB2 overexpression (Fig. 5d). We injected mice that were either WT, tOE, WT cKO (LATS1/2) or tOE cKO (LATS1/2) with tamoxifen for 7 days before transient ERBB2 induction (Fig. 5d). Successful LATS1/2 deletion was validated by western blot for pYAP S112 (a target residue of LATS1/2), whereby pYAP/Total YAP ratio was halved in the WT cKO and tOE cKO groups (Extended Data Fig. 5b-c). Echocardiographic analysis revealed that unlike the tOE group, which undergoes robust functional improvement upon ERBB2 shut-off (i.e. redifferentiation), tOE-cKO mice lacking LATS1/2 in CMs experienced a sustained and dramatic reduction in ejection fraction, fractional shortening, relative stroke volume and LV volume (Fig. 5e-f). WT cKO mice also experienced a decline in function, but to a lesser extent (Fig. 5e-f). This pattern between the groups was paralleled by the presence of dedifferentiation phenotypes at the Rediff timepoint in the cKO groups, such as disassembled sarcomeres, Nestin positive CMs, and pH3 positive CMs, which were totally absent in the WT and tOE groups Fig. 5g-h). This is consistent with other studies that show the role of the Hippo pathway in maintaining CM differentiation and maturity^8,40^. In summary, our findings suggest that under basal conditions, LATS1/2 is able to maintain CM identity but can also be activated as a countering force in response to ERBB2 signalling. In doing so it establishes a new equilibrium whereby CMs can proliferate, but are able to redifferentiate upon signal cessation. If however, LATS1/2 is disabled in combination with ERBB2 activation, dedifferentiation is maintained even after ERBB2 is shut off and redifferentiation is prevented (Fig. 5i). Hence, LATS1/2 is required for CM redifferentiation.

## Discussion

Physiologically, CMs that readily proliferate, contract weakly, such as those of adult zebrafish or neonatal mammals. Conversely, CMs that do not readily proliferate, such as those found in adult mammals contract strongly. This appears to be the evolutionary trade-off mammals made to meet the high oxygen demands of their bodies^41^. It therefore stands to reason that during mammalian heart regeneration, dedifferentiated CMs resembling those of neonates, lose some of their contractile ability, and therefore must undergo a return journey; redifferentiation, in order to recover their baseline function. Although there is growing appreciation of this process, it remains poorly understood^1^. Many studies have been published to date that claim heart regeneration by dedifferentiating CMs through an irreversible intervention, creating a paradox of regeneration without redifferentiation^8–15^. As awareness of this paradox has grown, more studies have begun to report the deleterious and lethal effects of persistent dedifferentiation^2–6^ and there has been a subsequent focus on transient interventions for heart regeneration^5,42–44^. It has been suggested that the process of dedifferentiation is reversible, based on the return to normal levels of individual markers after transient dedifferentiation in mouse^5^ or pig CMs^42^. Indeed, we too see many such markers return to normal, in line with declining ERBB2 levels and restoration of contractility. However, our high-throughput screens revealed a more complex picture, whereby a significant amount of genes and proteins remain or become differentially expressed at the Rediff timepoint, indicating that transient dedifferentiation of CMs may be less reversible than originally thought. Understanding the consequences of what does and does not return to normal will be key in translating regenerative medicine research to the clinic.

We found that hearts that underwent a dedifferentiation-redifferentiation cycle appeared rejuvenated, in that they had restored function, but showed lingering hallmarks of dedifferentiation, including remnants of cell cycle activity, EMT-like features and lower oxphos metabolism markers. We asked whether this would be beneficial against future ischaemic insult, and found that such hearts were robustly protected, with preserved contractile function and reduced scarring. We speculate several possible non-exclusive mechanisms for this effect. One is that the lingering dedifferentiation features, particularly those relating to reduced oxphos metabolism, are more accommodating to hypoxic conditions resulting from ischaemic injury^44,45^. Another is that the greater blood vessel density allows the myocardium to draw oxygen supplies from a wider source, reducing the impact of arterial occlusion. Another intriguing possibility is that the dramatic reduction in Cx43 expression blocks the spread of necrotic signalling molecules originating from the initial ischaemic event, which are normally mediated by Cx43 gap junctions^46^ and hemichannels^47,48^. This appears to mimic the cardioprotective effects of Cx43 blocking peptides^49^. The significance of this finding is underscored by the medical community’s interest in prophylactic therapies for ischaemic damage such as ischaemic preconditioning, which despite being first reported over 30 years ago, has struggled to reach the clinic^50^. As the promises of cardiac regeneration are realised and the idea of transient mitogenic signalling in human CMs gains traction, so too might the prophylactic use of these therapies to confer cardioprotection.

During the dedifferentiation phase, we noted a profound and wide-ranging induction of negative feedback regulators, representing a robust resistance to mitogenic signalling, which subsided in step with ERBB2 levels. We also showed that this pattern is mirrored in wildtype mice from birth to adulthood, suggesting that negative feedback signalling in neonates may be important for CM differentiation and maturation. This finding may prove useful for researchers generating iPSC derived CMs, whose efforts have been hampered by insufficient maturation^51^. Negative feedback signalling may also explain the paradox posed above, whereby irreversible dedifferentiation signals are sufficiently countered to stabilise heart function. This is consistent with a study whereby overexpression of a cocktail of four cell-cycle activators caused a burst of CM proliferation which diminished rapidly after one week, in response to their increased proteasomal degradation^52^. A similar trajectory was seen after CM-specific deletion of the Hippo pathway component *Salv*, which caused an initial increase in cell cycle activity, before waning in the weeks thereafter^13^. Nonetheless, the role of negative feedback signalling in regeneration as the driver of redifferentiation has not been previously established or studied. Here, we showed that inhibiting or disabling negative feedback signalling via the major kinases LATS1/2 augmented dedifferentiation *in vitro* and sustained dedifferentiation *in vivo*, highlighting the necessity of LATS1/2 Hippo signalling for successful redifferentiation. In contrast to the widely accepted notion of CM terminal differentiation in the adult mammalian heart, our study adds to the growing body of evidence that Hippo signalling is actively required to maintain CMs in a differentiated state^8,40^. Although we focussed our study on LATS1/2 and the Hippo pathway, we speculate that other negative feedback regulators are also important in redifferentiation, such as those that target ERK1/2, AKT and ERBB2 itself. Dusp5^53^ and Dusp6^54^, which dephosphorylate activated ERK1/2, are attractive candidates as negative feedback regulators in redifferentiation. We demonstrate their upregulation in dedifferentiated mouse hearts, and *Dusp6* upregulation in actively cycling zebrafish CMs. This is consistent with studies showing that their inhibition can further promote CM proliferation in zebrafish, mice and rats^16,53–55^.

Our study focussed on the redifferentiation of the heart with an emphasis on CMs, however it is clear from histological analysis that forced CM dedifferentiation affects many cell types, and their relative composition in the heart. It will be equally important in the future to characterise all cells of the redifferentiated heart, and the consequences of any irreversible changes. A lot of damage was incurred to the field of cardiac regenerative medicine when researchers rushed to trial stem cell-based cardiac regenerative therapies in humans, which inevitably failed owing to our poor understanding of the underlying mechanisms of regeneration^56,57^. Similarly, at a time when our understanding of the first phase of cardiac regeneration i.e. CM dedifferentiation is so advanced, it will be important to understand the second phase, redifferentiation, in sufficient detail before progressing to the clinic again.

## Methods

### Mouse Experiments

Mouse experiments were approved by the Animal Care and Use Committee of the Weizmann Institute of Science (approval no. 13240419-3 and 05850721-2) and the study is compliant with all of the relevant ethical regulations regarding animal research. Doxycycline-inducible CM-specific caERBB2 overexpression was achieved by crossing the TetRE-caERBB2 mouse line^58^ (a B6SJLF1/J transgenic line in which the activated form of rat *c-neu*/erbB2 is placed under the control of a minimal CMV promoter and the tetracycline-responsive element) with an αMHC–tTA^59^, which expresses the tetracycline-responsive transcriptional activator (tTA) under the control of the human alpha myosin heavy chain promoter. The αMHC–tTA was originally created in the FVB background and was subsequently backcrossed into the C57BL/6 background. Doxycycline hyclate-supplemented special diet (Dox, Harlan Laboratories, TD.02503, 625mg/kg) was fed to the mice *ad libitum* to repress transgene expression. To track the cardiac muscle cell lineage, we intercrossed αMHC–Cre to ROSA26–tdTomato mice as previously described^18^.

For the Lats1/2 cKO mice, inducible-Cre mice (αMHC-MerCreMer; Jackson, 005657) were crossed with Lats1/2 flox^8,40^ mice and then with αMHC-tTA all in one mouse, while TetRE-caERBB2 were crossed with Lats1/2 flox mice and ROSA26::tdTomato all in another mouse. These mice were crossed and mice inheriting all transgenes were OE-cKO and those lacking αMHC-tTA were WT-cKO. A list of genotyping primers for the transgenes used in this study are detailed in Supplementary Table 1.

### Myocardial infarction

Myocardial infarction in adult mice was achieved by ligation of the left anterior descending coronary artery as previously described^3,18^. Briefly, mice were anaesthetised with isofluorane (Abbott Laboratories) and artificially ventilated following tracheal intubation. Lateral thoracotomy at the fourth intercostal space was performed by blunt dissection of the intercostal muscles following skin incision. Following LAD ligation, thoracic wall was closed and the incision sealed with Histoacryl® (B. Braun Melsungen AG) and the mice were administered buprenorphine (0.066 mg/kg) by subcutaneous injection as an analgesic and warmed until recovery.

### Echocardiography parameters

Cardiac function was evaluated by transthoracic echocardiography performed on isofluorane (2-3%, Abbot Laboratories) sedated mice using the Vevo3100 (VisualSonics), keeping the heart rate between 340 and 510bpm. Ejection fraction, systolic volume, diastolic volume, and stroke volume were calculated from the long axis, whilst measurements for the left ventricular anterior wall and left ventricular posterior wall were calculated from the short axis, representing the papillary plane thickness. Parameters were calculated using Vevo Lab 3.2.6 software (VisualSonics). Mice with a 2-week ejection fraction value lower than 40% or higher than 90% of their baseline value were excluded from experiments in Fig. 1 and Fig. 5 as the injury was determined as either too severe for ERBB2 to induce regeneration or insufficient to be considered a true injury. No mice were excluded for experiments in Fig. 3 since they could not be distinguished from genuine cardioprotection or injury sensitivity. All analysis performed was blinded except for during ERBB2 overexpression since the phenotype presents robustly in an identifiable manner.

### RNAseq

RNA-seq was performed as previously described^18^. Briefly, RNA was extracted and purified from PBS-perfused, liquid N_2_-frozen, powdered whole heart tissue using an miRNeasy kit (Qiagen, 217004) followed by library preparation (G-INCPM, Weizmann Institute of Science). Sequencing libraries were constructed and read using an Illumina HiSeq 2500 machine, using the Single-Read 60 protocol (v4). Poly-A/T stretches and Illumina adapters were trimmed from the reads using cutadapt^60^ and reads that were shorter than 30 nucleotides were discarded. Reads were aligned to the mm10 mouse genome (STAR)^61^, supplied with gene annotations downloaded from Ensembl (and with EndToEnd option and outFilterMismatchNoverLmax was set to 0.04). Expression levels for each gene were quantified using HTseq-count^62^. Differential expression analysis was performed using DESeq2 (1.6.3)^63^ with the betaPrior, cooksCutoff and independentFiltering parameters set to False. Raw *P* values were adjusted for multiple testing (Benjamini and Hochberg).

Principal component analysis was performed on the DESeq2 variance stabilizing transformed values, of the 1,000 most variable genes (using the R Stats package). To create the dendrogram, hierarchical clustering was performed on the DESeq2 variance stabilizing transformed values of the 1,000 most variable genes. Pearson’s correlation was used as a distance metric with the Ward algorithm.

### Proteomics

Proteomics was performed as previously described^18^. Briefly, protein was extracted from the powder (see the ‘RNA-seq’ section) by homogenisation in SDT buffer supplemented with phosphatase and protease inhibitors. Samples were subjected to tryptic digestion followed by liquid chromatography separations as previously described^64^. Peptides were detected using Mass spectrometry (Q Exactive Plus, Thermo Scientific) and the raw data were imported into the Expressionist software (Genedata, version 9.1.3) and processed as described previously^65^. Data were searched against the mouse protein database downloaded from UniprotKB (http://www.uniprot.org/). Peptide identifications were imported back to Expressionist to annotate the identified peaks. Quantification of protein abundance from the peptide data was performed using an in-house script^65^, obtained by summing the three most intense unique peptides per protein. A Student’s t-test, after logarithmic transformation, was used to identify significant differences across the biological replicates. Fold changes were calculated based on the ratio of the arithmetic means of the experimental groups. The PCA and hierarchical clustering were calculated by their respective functions in Perseus.

### Bioinformatics

The Ingenuity Pathway Analysis tool (Qiagen Inc.) was used for the analysis of the RNA-seq and proteomics data using differential expression thresholds (as stated in the figure legend). Heat maps were constructed using Morpheus (https://software.broadinstitute.org/morpheus). GO term enrichment for Rediff RNA-seq data was performed using PANTHER software^66^(http://www.pantherdb.org) testing for overrepresentation with a significance threshold of 0.05 for the adjusted *q*-value.

### RT-qPCR

RNA from whole hearts was isolated using a miRNeasy RNA extraction kit (Qiagen, 217004) according to the manufacturer’s instructions. A High capacity cDNA reverse transcription kit (Applied Biosystems, 4374966) was used to reverse transcribe 2µg of purified RNA according to the manufacturer’s instructions. The quantitative PCR reactions were performed using a Fast SYBR Green PCR master mix (ThermoFischer Scientific, 4385614). Expression FC values were calculated using the 2^−ΔΔCt^ method using StepOne (ThermoFischer Scientific, v2.3). The oligonucleotide sequences used for RT-qPCR analysis performed in this study are listed in Supplementary Table 2.

### Western Blot Analysis

Total tissue lysates were isolated using RIPA buffer supplemented with 1:100 protease (Sigma, P8340) and 1:100 phosphatase inhibitor cocktails (Sigma, P5726 and P0044). Protein concentrations were quantified by BCA assay (Thermo Fisher, 23225). The lysates were separated on 4-20% Mini-PROTEAN® TGX Stain-Free(tm) gels (BIO-RAD) and wet transferred onto 0.44µm PVDF membranes. The membranes were blocked and probed with primary antibodies according to manufacturer’s instructions (detailed list in Supplementary Table 3). The results were visualised and analysed using Image lab software (v6.1)

### Immunofluorescence

#### Heart sections

Hearts were briefly perfused *in situ* with ice-cold PBS for 30s and then by 4% paraformaldehyde (PFA) for 1 min, followed by overnight incubation shaking at 4°C. The samples were then embedded in paraffin and cut into 5μm thick sections. Slides were deparaffinised and heat-mediated antigen retrieval was performed using EDTA or citrate buffer, followed by 5 minutes of permeabilisation (0.5% Triton X-100), 1h of blocking and addition of primary antibodies in blocking solution (3% BSA, 10% heat-inactivate Horse/Goat serum (Biological Industries: 04-124-1A, 04-009-1A) in 0.1% Triton X-100) for overnight incubation at 4°C (detailed list in Supplementary Table 4). A hydrophobic pen was used to draw a perimeter around the sections, promoting equal coverage of the antibody solution across all sections. Sides underwent 3 cycles of 10 minute PBS washes followed by the application of secondary antibodies (Jackson, Abcam) with DAPI for 40 min at room temperature. The slides were washed and mounted with Immu-Mount (Thermo Scientific, 9990402) and imaged on a Nikon Eclipse Ti2 microscope.

#### P7 cardiac cultures

Cells were fixed with 4% paraformaldehyde for 12 min, followed by permeabilisation with 0.5% Triton X-100 for 5 min and blocking with 5% bovine serum albumin and 0.1% Triton X-100 for 1hr. Primary antibody solution was prepared in blocking solution (detailed list in Supplementary Table 3) and incubated on an orbital shaker for 3hr at room temperature. The cultures subsequently underwent 3 cycles of 10 min PBS washes and incubated for 40 min with the appropriate secondary antibodies (Abcam or Jackson) and DAPI. After an additional 3 cycles of 10 min PBS washes, the cultures were imaged on a Nikon Eclipse Ti2 microscope.

### Respiration and Glycolysis

Measurements of live cellular respiration and ECAR were measured using the Seahorse XFe96 Analyzer (Agilent) according to the manufacturer’s instructions. P7 hearts were digested as described below, with an additional step to purify the culture for CMs (Miltenyi Biotech, 130-100-825). ECAR was measured using the Glyco Stress Test (Agilent, 103020-100) and OCR was measured using the Mito Stress Test (Agilent, 103015-100). Data were normalised to cTnT fluorescent staining intensity of Seahorse plate wells following the experimental runs in the Analyzer.

### Optical Mapping

Optical mapping was performed as previously described^67,68^. Briefly, hearts were extracted from anaesthetised mice and washed in oxygenated (95%) 37°C Tyrode’s solution. Once cannulated via the aorta, the heart was perfused with Tyrode’s solution with 20 mM 2,3-Butanedione monoxime. Before measurement, the perfusate was spiked with 1ml of 100μM CytoVolt1(Di-4-ANBDQBS) and 200μM blebbistatin in Tyrode solution over the course of 2 minutes. Optical mapping was performed using a high speed CCD based technique (Evolve-512 Delta, Photometrics) with the hearts paced at a constant rate of 200CL. The X-Cite Turbo LED-system served as a light source. ElectroMap software was used for analysis of optical signals. The data were viewed initially as dynamic displays showing the propagation of the activation wavefronts. Activation maps were then constructed by measuring the timing of electrical activation at each image pixel (timing of the maximum dF/dt).

### Scar Analysis

Scar quantification was performed in a blinded manner, based on Sirius red staining of serial cardiac sections spanning the entire heart. In each section, the area of fibrotic tissue as a percentage of the area of the left ventricle was measured using an in-house script within ImageJ (1.52q). This was then averaged across all sections. For scar class, each section was graded as having a transmural scar (i.e. scar spanning the entire width of the left ventricle at any point), non-transmural scar or no scar. The number of sections for each category was expressed as a percentage of total sections on the slide.

### Re-analysis of Published Data Sets

Two different bulk RNAseq data sets were reanalysed in this study. The first, from O’Meara et al. contained data from whole heart lysates from P0, P4, P7 and adult mice (*n* = 2 per group) and was obtained in a completely analysed form with FPKM values^36^. The second, from Quaife-Ryan et al. was generated by separating myocyte and non-myocyte fractions of dissociated hearts, before FACS sorting the non-myocyte fractions into endothelial cells, fibroblasts and immune cells^37^. Raw data was downloaded from the GEO database (GSE95755).A count matrix for each gene was quantified using HTseq-count^62^ on union mode. and differential expression analysis was performed with EdgeR (v3.2.4)^69^.

### Zebrafish ISH and FISH

All animal experiments were conducted under the guidelines of the animal welfare committee of the Royal Netherlands Academy of Arts and Sciences (KNAW). Adult zebrafish (Danio Rerio) were maintained and embryos raised and staged as previously described (PMID: 31510859, Westerfield, M. (2000) The Zebrafish Book. A Guide for the Laboratory Use of Zebrafish (Danio rerio), 4th Edition. University of Oregon Press, Eugene.) Tupfel longfin (wild type) between 6months and 1 year were subjected to cryoinjury as previously described^70^. At 7 days post injury, hearts were extracted and fixed in 4% PFA overnight at 4oC. Hearts were then washed 3x in 4% sucrose/PBS and allowed to equilibrate in 30% sucrose/PBS at 4°C for 5 hours. Hearts were then suspended in Tissue Freezing Medium (Leica) and frozen with dry ice. Blocks were cryosectioned at 10um at 20°C. Slides were subjected to the in situ hybridization protocol as previously described^19^ with the exception of using fast red (Sigma) to develop the signal. *In situ* probes were made by PCR with T7 RNA polymerase as previously described^71^ using the following primers listed in Supplementary Table 5. Once the *in situ* stain was developed, slides were immediately washed in PBST and an antigen retrieval step with 10mM Sodium Citrate at 84°C for 15 mins, permeabilised with 0.1% collagenase II in PBS with 0.1% Tween20 and blocked for at least 30mins in PBS containing 10% FCS, 1% DMSO, 0.1% Tween. Primary antibodies (PCNA, Dako #M0879 1:800 dilution; tropomyosin, Sigma #T9283, 1:500 dilution) were diluted in blocking buffer overnight at 4°C. Followed by 3x washes in PBS with 0.1% Tween20, incubation with Alexa Fluor secondary antibodies at 1:500 dilution and DAPI. Slides were imaged on an VS200 slide scanner (Olympus) or LSM900 confocal (Zeiss). Due to anti-Tropomyosin primary antibody cross-reactivity with PCNA, we performed post-acquisition processing of images to dampen the artefactual Tropomyosin signal in overlapping regions i.e. PCNA positive nuclei, to a uniform intensity matching that of the cytoplasmic regions. This procedure was performed uniformly in all acquired images, and was only used for illustrative purposes. Quantification was performed on un-processed images.

### CM Isolation and culture

CM Isolation and culture was performed as previously described^18^. Briefly, cardiac cultures were isolated from P7 mice using a neonatal dissociation kit (Miltenyi Biotec,130-098-373) and the gentleMACS homogeniser, according to the manufacturer’s instructions, and cultured in ‘complete medium’ at 37°C in 5% CO_2_ for 24h, before being replaced by ‘starvation media’ for an additional 48h. To inhibit LATS1/2, the LATS1/2 inhibitor TRULI^39^ was used at a final concentration of 10uM in the ‘starvation’ media. For LATS1/2 conditional knockout, pups were administered a daily subcutaneous injection of 10-20ul Tamoxifen (10mg/ml) from P3 to P7. Following isolation, cells were cultured in the media described above supplemented with 10uM Tamoxifen (Sigma, T5648).

### Time-lapse movie and single-cell tracking

P7 WT, OE, WT cKO or OE cKO cardiac cultures tagged with tdTomato fluorescent protein were cultured as described above. After 24h, the medium was replaced and time-lapse imaging began for 48h, with images acquired every 10m. Proliferation was expressed as a percentage of full CM cell division events divided by number of CMs per field by the last frame.

### Statistics and reproducibility

All experiments were carried out with n ≥ 3 biological replicates. Experimental groups were balanced for animal age, sex and weight. The animals were genotyped before the experiment, caged together and treated in the same way. Statistical analyses were carried out using the Prism software (version 6.0.1). When comparing between two conditions, data were analysed using a two-tailed Student’s t-test (except for Fig. 1g, where a one-tailed t-test was used). When comparing more than two conditions, we employed an ANOVA analysis with multiple comparisons. In bar plots comprised of colour coded dots (such as in Fig. 1c), the dots represent data points derived from different biological repeats to demonstrate the distribution of data across different experiments. The statistical analysis is derived from the biological repeats of an experiment. Measurements are reported as the mean and the error bars denote the s.e.m. throughout the study. The threshold for statistical significance was considered as \**p* ≤ 0.05, \*\**p* < 0.01, \*\*\**p* < 0.001, \*\*\*\**p* < 0.0001, and ns, not significant, as stated in the figure legend.

## Acknowledgements

This study has been supported by grants to E.T. from the European Research Council (ERC StG grant no. 281289, CM turnover, and ERC AdG grant no. 788194, CardHeal), ERA-CVD CARDIO-PRO, EU Horizon 2020 research and innovation programme REANIMA, the U.S.–Israel Binational Science Foundation (BSF), the Israel Science Foundation (ISF), Foundation Leducq Transatlantic Network of Excellence and Minerva foundation, with funding from the Federal German Ministry for Education and Research. We thank the Benoziyo Endowment Fund for the Advancement of Science, Head of the Yad Abraham Research Center for Cancer Diagnostics and Therapy, Zuckerman STEM Leadership Program, Dr. Dvora and Haim Teitelbaum Endowment Fund and Daniel S. Shapiro Cardiovascular Research Fund. This work was supported by grants from the EMBO Long Term Fellowship ALTF1129-2015, HFSPO Fellowship (LT001404/2017-L) and a NWO-ZonMW Veni grant (016.186.017-3) (P.D.N), the Netherlands Cardiovascular Research Initiative: An initiative with support of the Dutch Heart Foundation and Hartekind, CVON2019-002 OUTREACH (J.B.) and the National Institutes of Health grant T32GM007739 (N.K.). We thank Ofra Golani for assistance with image processing. Illustrations in figures were created with BioRender.com.

## Author contributions

A.S. and E.T. conceived and designed the experiments. A.S., with help from Z.P. and A.A. carried out most of the experiments and analysed the data. M.G. performed the optical mapping studies. K.B.U. helped with the animal studies, RNA-seq and proteomic preparation. P.D.N. and J.B. conducted zebrafish related experiments. D.K. and D.L. performed the LAD-ligation and echocardiography acquisition. J.E. helped with various IHC experiments. Y.D. performed scar analysis. G.F. performed RNA-seq analysis. A.Savidor and Y.L. performed proteomic analysis. S.M., L.Z. and D.E.P. performed various supporting experiments that guided the project and N.K. provided the LATS inhibitor. A.A. and K.B.U. contributed to the planning of the project. E.T. supervised the entire project and A.S. and E.T. wrote the manuscript with contributions from all of the authors.

---

**Extended Data Fig. 1.**
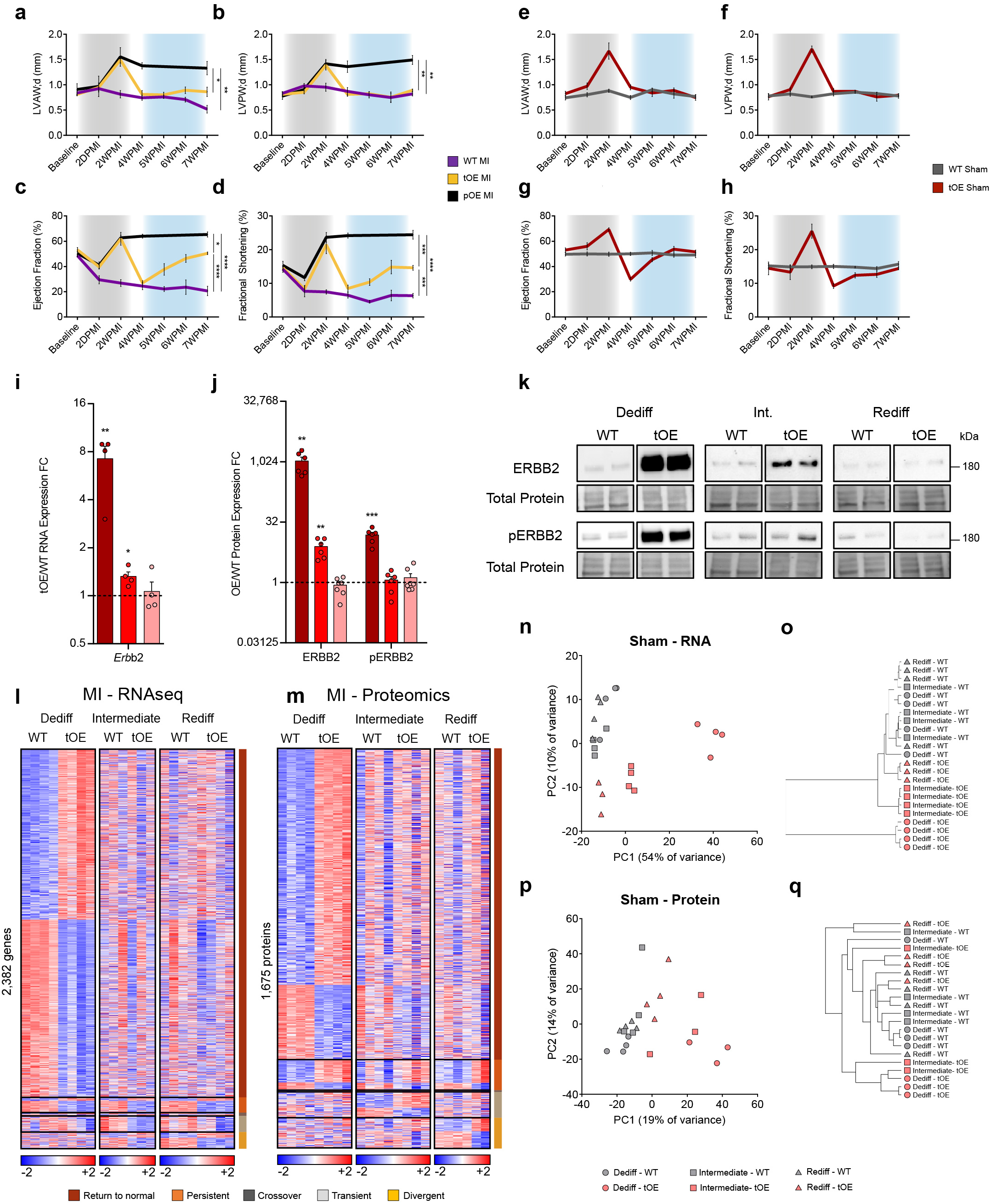
Functional recovery after transient ERBB2 signalling occurs despite incomplete reversal and development of differentially expressed genes and proteins. **a-h**, Left Ventricular Anterior Diastolic Wall thickness (LVAW;d) **(a**,**e)**, Left Ventricular Posterior Diastolic Wall thickness (LVPW;d) **(b**,**f)**, Ejection Fraction **(c**,**g)**, and long-axis Fractional Shortening **(d**,**h)**, of WT Ml, tOE Ml, pOE Ml, WT Sham and tOE Sham mice, measured by echocardiography. Gray shaded area represents time period of caERBB2 activation (dedifferentiation). Blue shaded area represents the redifferentiation phase. Overall, *n* = 3 - 8 per group. **i**, Sham tOE/WT RNA expression fold change (FC) of *Erbb2*. All values are normalised to their in-time point average WT value (black dashed line). All groups had minimum n = 3. **j** Western blot quantification of tOE/WT FC of ERBB2 and pERBB2 (Tyr-1248) protein. All values are normalised to their in-time point average WT value (black dashed line). *n* = 4 - 8 per group. **k** Representative western blot images of data from (j). **1-m**, Heatmaps of differentially expressed genes (I) and proteins **(m)** from Ml-injured samples, compiled as described in Fig. 1g-h. **n-q** Principal component analysis and dendrograms of sham from sham RNAseq **(n**,**o)** and sham proteomics groups **(p**,**q)**. In all panels numerical data are presented as mean ± SEM; statistical significance was calculated using one-way ANOVA with Sidak’s multiple comparison test at the 7WPMI time point in **a-h**, two-tailed unpaired Student’s t-test between tOE and WT of each time point in i and **j**. *\*p* 0.05, *\*\*p* < 0.01, *\*\*\*p* < 0.001, *\*\*\*\*p* < 0.0001.

**Extended Data Fig. 2.**
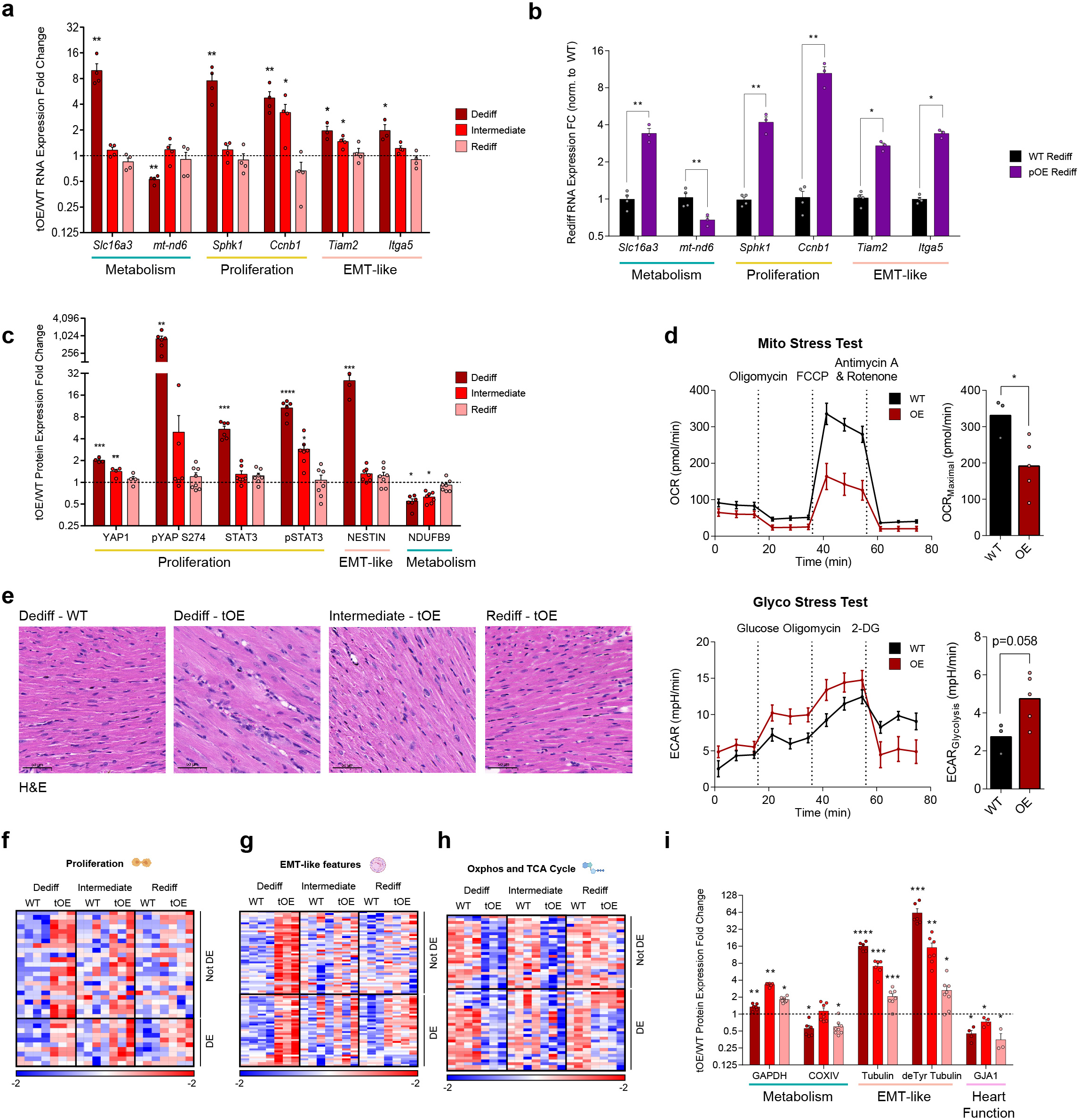
Dedifferentiated phenotypes are largely reversed in functionally redifferentiated hearts. **a-b** tOE/WT at all timepoints **(a)** and pOE/WT at Rediff **(b)** RNA expression fold change of genes that returned to normal, involved in metabolism, proliferation and EMT-like features, determined by RT-qPCR. All values are normalised to their in-time point average WT value (black dashed line). *n* = 3 - 4 per group. **c** Western blot quantification for data in Fig. 2b in order to validate the ‘return to normal’ behaviour of proteins involved in proliferation, EMT-like features and metabolism. **d** Metabolic analysis of cultured P7 WT *(n* = 3) and OE *(n* = 5) CMs using an XFe96 Seahorse analyser. Upper left panel shows the OCR (oxygen consumption rate) during the Cell Mito Stress Test and lower left panel shows the ECAR (extracellular acidification rate (glycolysis proxy)) during the Glycolysis Stress Test. Maximal respiration/OCR and Glycolysis are represented in the upper right and lower right panels respectively. e H&E stained histological sections of WT Dediff and tOE Dediff, Int. and Rediff hearts. Images were acquired in the remote zones of Ml injured hearts as a proxy for sham injury. Scale bars = 50µm. **f-h** Heat maps based on log’°transformed intensity values for Sham proteomics. Rows represent proteins. Columns represent each biological sample. Colour bars represent z-score for each row across all timepoints. **i** Western blot quantification for data in Fig. 2g in order to validate the proteins involved in metabolism, cytoskeletal signalling and heart function that remain DE at Rediff. In all panels numerical data are presented as mean ± SEM; statistical significance was calculated using two-tailed unpaired Student’s t-test in **(a)-(d)** and (i) between the in-time point WT and tOE values. *\*p* 0.05, *\*\*p* < 0.01, *\*\*\*p* < 0.001, *\*\*\*\*p* < 0.0001.

**Extended Data Fig. 3.**
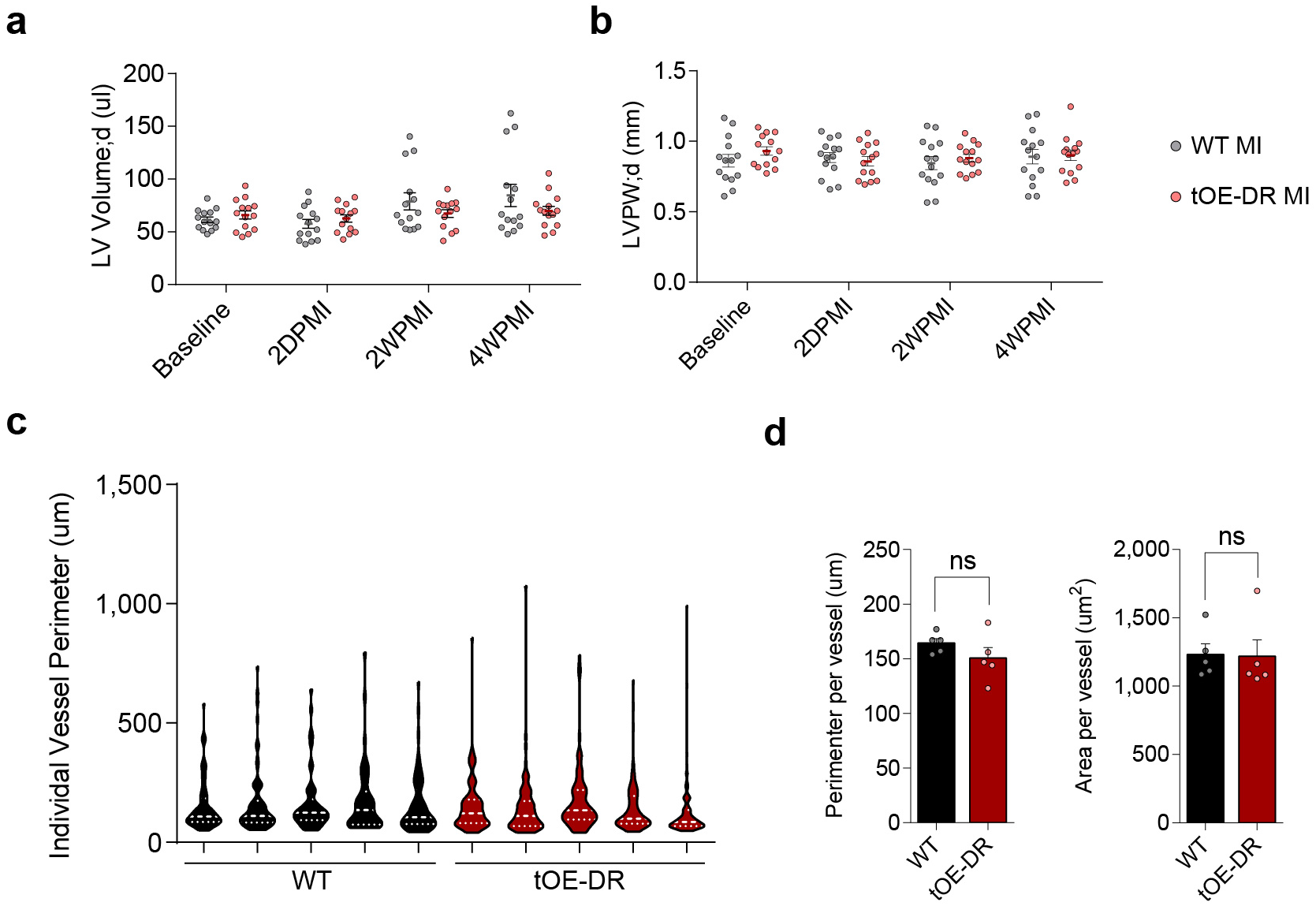
Dedifferentiation-Redifferentiation cycle does not affect ventricular dimensions after injury or baseline vessel size distribution. **a-b** Left ventricular diastolic volume (LV;d) **(a)** and Left Ventricular Posterior Diastolic Wall thickness (LVPW;d) of WT and tOE-DR, measured by echocardiography. *n* = 14 for each group. **c-d** Violin plot of vessel perimeter for each biological replicate quantified in Fig. 3k-l **(c)**, average perimeter (left panel) and area (right panel) per vessel **(d)** for WT *(n* = 278 vessels) and tOE-DR *(n* = 302) across 5 biological replicates per group. Numerical data in **(a), (b)** and **(d)** are presented as mean ± SEM, and as median (thick dashed white line) ± interquartile range (thin dashed white line) in **(c);** statistical significance was calculated using two-tailed unpaired Student’s t-test in **(a), (b)** and **(d)** between the in-time point (where applicable) WT and tOE-DR values. *\*p* 0.05, *\*\*p* < 0.01, *\*\*\*p* < 0.001, *\*\*\*\*p* < 0.0001

**Extended Data Fig. 4.**
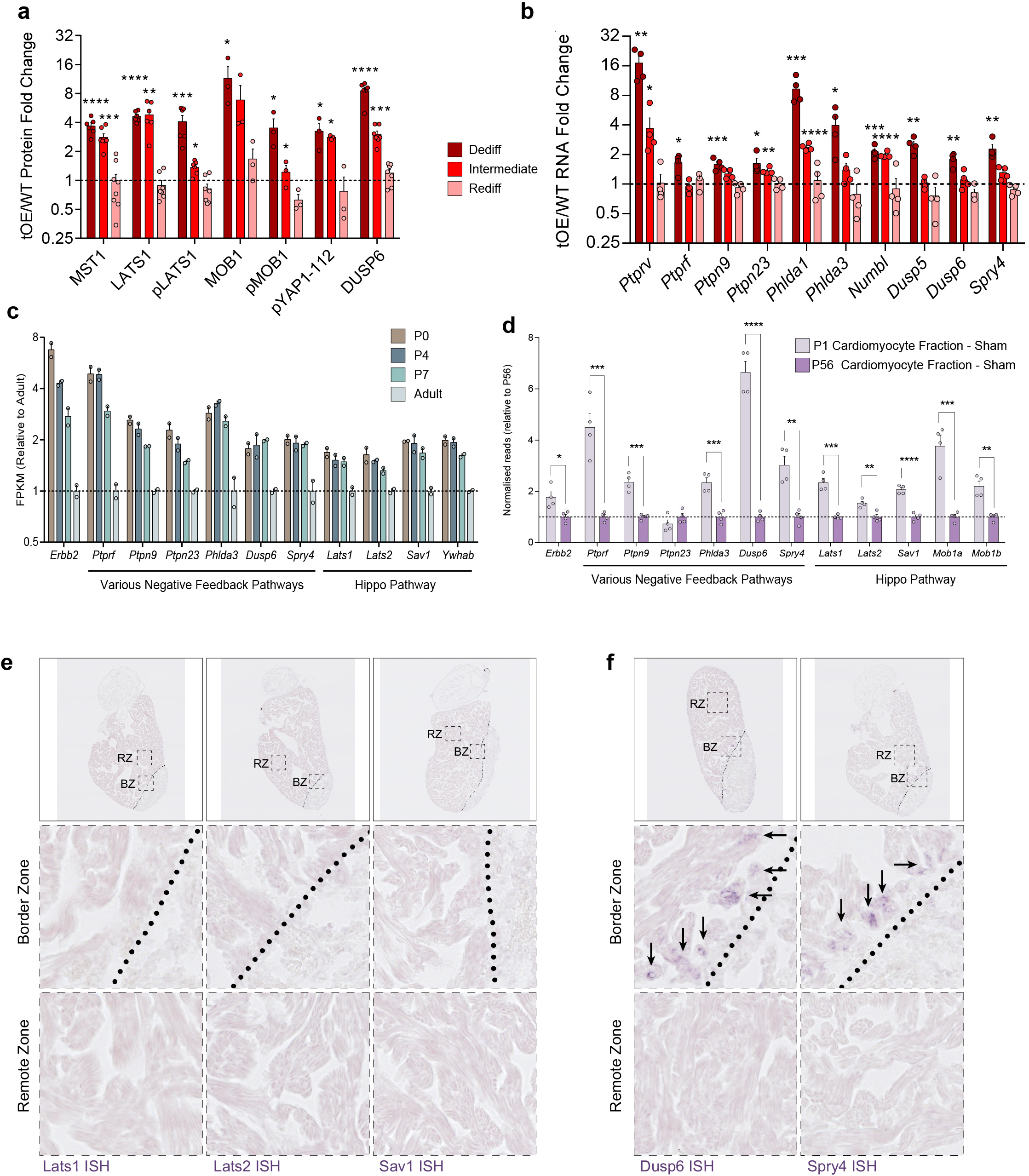
ERBB2 signalling promotes a multi-faceted negative feedback response. **a** Western blot quantification of whole-heart lysates for Hippo pathway components and Dusp6 for WT and tOE adult heart lysates for each time point. *n* = 4 - 8 per group. **b** qRT-PCR from whole-heart lysates for negative feedback (NF) regulators. **c** RNAseq FPKM values for NF regulators from ventricular lysate of various ages. Data re-analysed from O’meara et al. 2015^36^. **d** RNAseq normalised counts for NF regulators from bulk-RNAseq of purified cardiomyocytes from sham injured P1 and P56 mice. Data re-analysed from Quaife-Ryan et al. 2017^37^. **e-f** *In situ* hybridisation for Hippo pathway genes *Lats1, Lats2* and *Sav1* **(e)** and ERK NF regulators *Dusp6* & *Spry4* **(f)** mRNA in 7DPI adult zebrafish hearts. RZ = Remote Zone. BZ = Border Zone. Zones are delineated by the black dotted lines. Middle and bottom panels show higher magnification images of the corresponding dashed black boxes in the top panel. Black arrows highlight the presence of detected mRNA. In all panels numerical data are presented as mean± SEM; statistical significance was calculated using two-tailed unpaired Student’s t-test in **(a), (b)** and **(d)** between the in-time point WT and tOE values. *\*p* 0.05, *\*\*p* < 0.01, *\*\*\*p* < 0.001, *\*\*\*\*p* < 0.0001.

**Extended Data Fig. 5.**
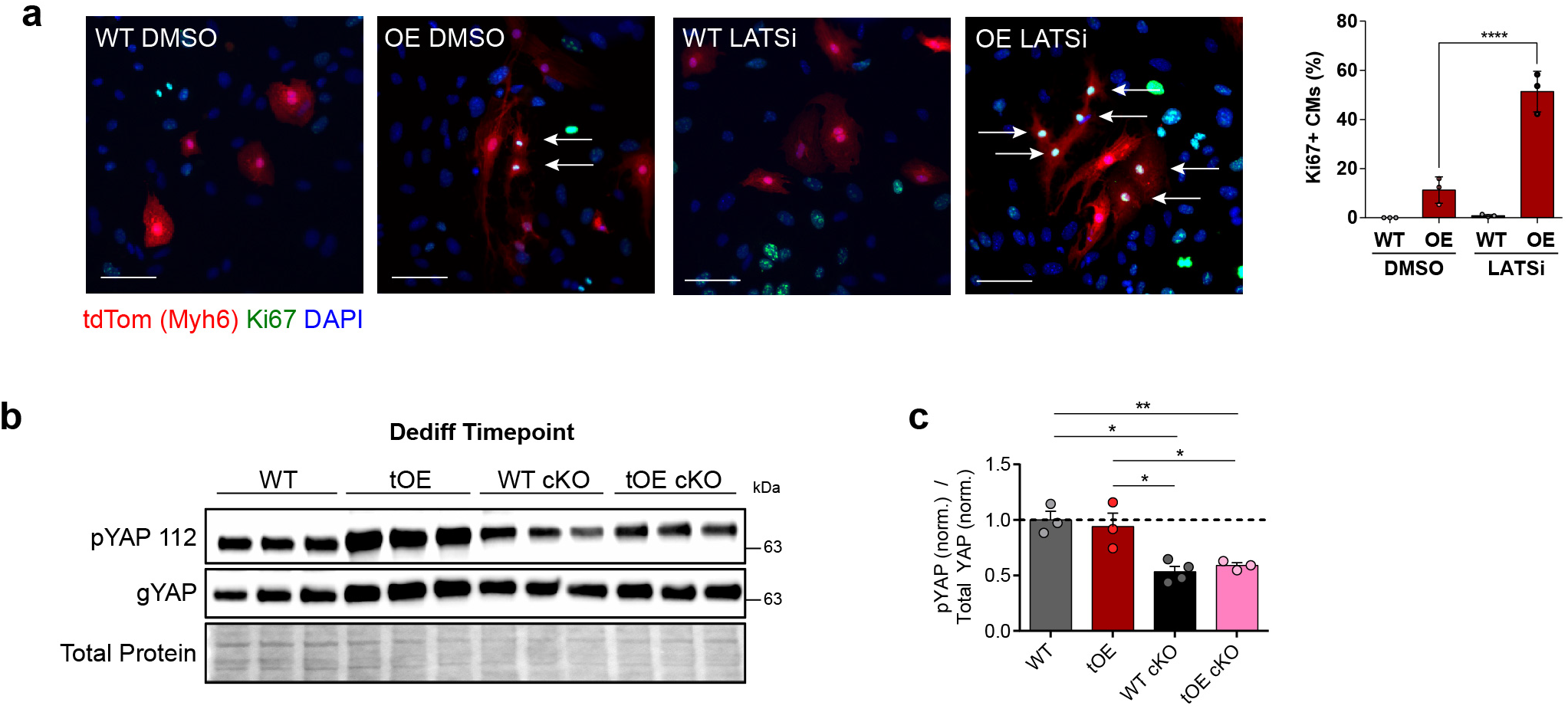
Disabling Hippo pathway prevents cardiomyocyte redifferentiation. **a** Representative immuno fluorescence images of Ki67 in LATSi or DMSO treated WT and OE P7 cardiac cultures, with full quantification. All groups had minimum *n* = 3. Scale bars = 50µm. **b-c** Representative western blot of whole-heart lysates for general Yap (gYAP) and pYAP S112 (a target residue of LATS1/2) from WT, tOE, WT LATS1/2 cKO and tOE LATS1/2 cKO mice **(b)**, with quantification, normalised to the average WT value **(c)**. All groups had minimum *n* = 3. In all panels numerical data are presented as mean ± SEM; statistical significance was calculated using a paired two-way ANOVA followed by Sidak’s test in **(a)** and a one-way ANOVA followed by Tukey’s test in **(c)**. *\*p* :=; 0.05, *\*\*p* < 0.01, *\*\*\*p* < 0.001, *\*\*\*\*p* < 0.0001.

## Supplementary Data Tables

**Table 1.**
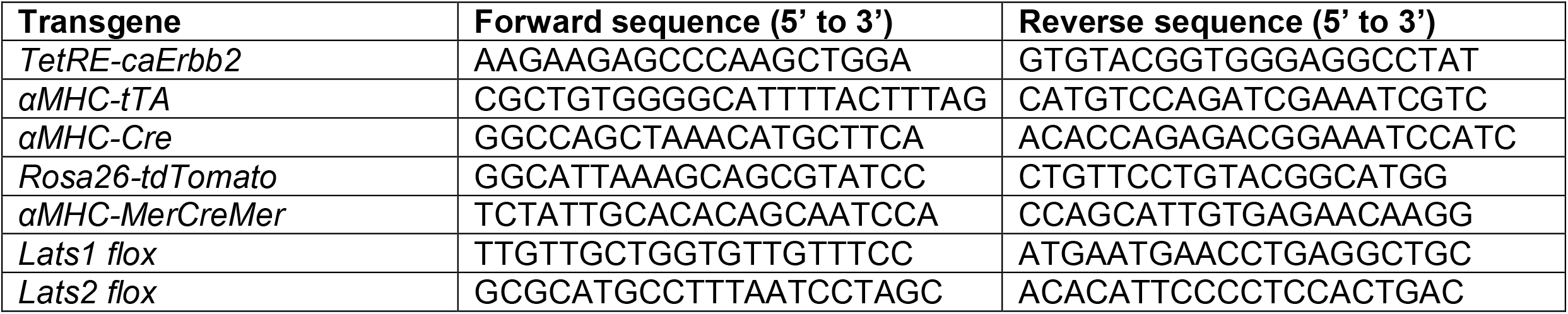
Genotyping Primers used for Transgenic Lines.

**Table 2.**
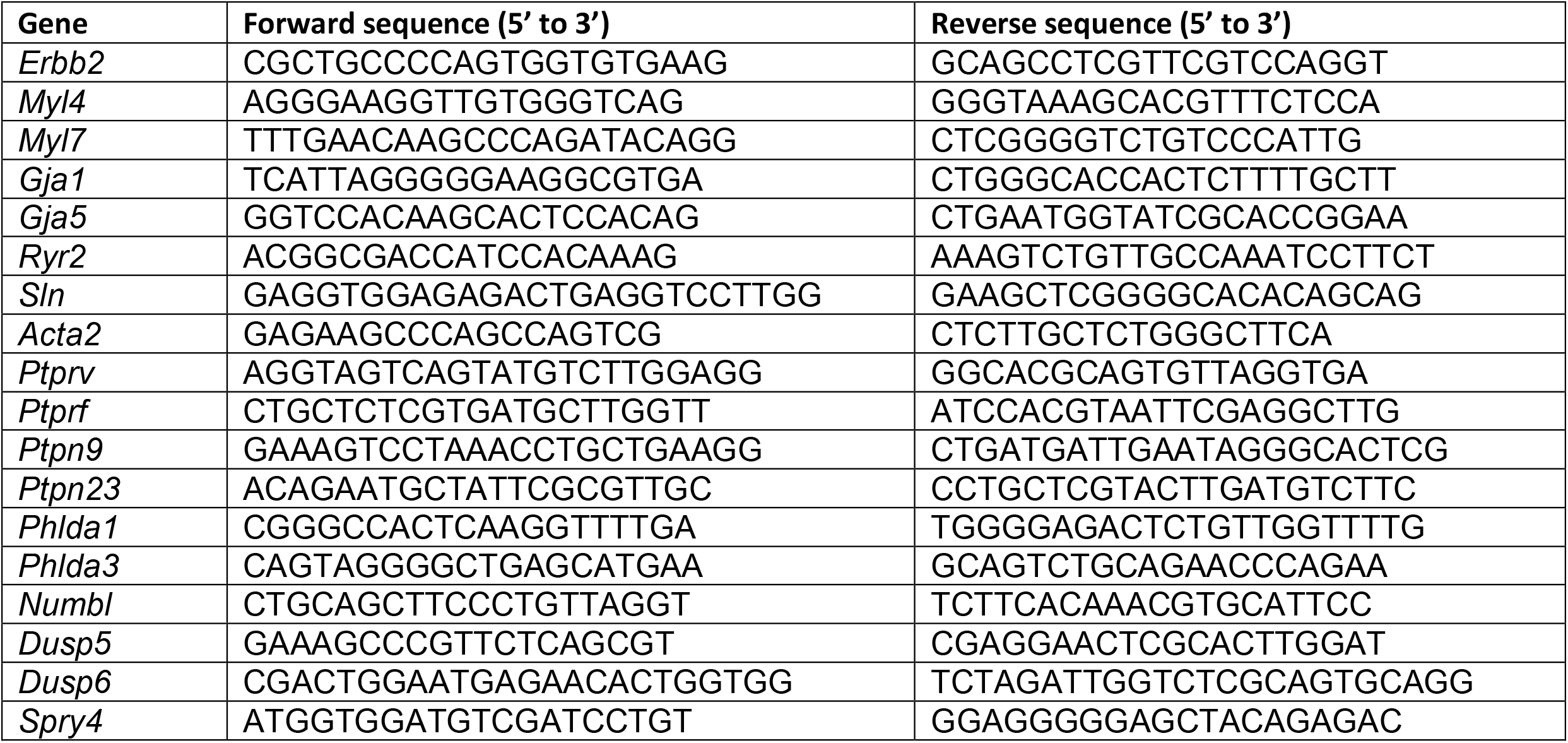
Primers for RT-qPCR.

**Table 3.** Antibodies used for Western Blots.

| Antibody against | Manufacturer | Cat. No. | Dilution |
| --- | --- | --- | --- |
| HER2/ERBB2 | Cell Signalling | #2165 | 1:1000 |
| pHER2/ERBB2 (Tyr1248) | Cell Signalling | #2247 | 1:1000 |
| pYAP (Ser274) | n/a | Custom Ab. | 1:1000 |
| YAP | Novus Biologicals | NB110-58358 | 1:1000 |
| STAT3 | Cell Signalling | 12640 | 1:1000 |
| pSTAT3 (Tyr705) | Cell Signalling | 9145 | 1:2000 |
| NESTIN | Abcam | ab11306 | 1:1000 |
| NDUFB9 | Abcam | ab188581 | 1:10000 |
| GAPDH | Sigma | PLA0125 | 1:2000 |
| COXIV | Abcam | ab16056 | 1:1000 |
| deTyrosinated Tubulin | Abcam | ab48389 | 1:1000 |
| TUBULIN | Cell Signaling | 3873 | 1:1000 |
| GJA1 | Abcam | ab11370 | 1:5000 |
| MST1 | Cell Signaling | 3682 | 1:1000 |
| LATS1 | Cell Signaling | 3477 | 1:1000 |
| pLATS1 (Ser909) | Cell Signaling | 9157 | 1:1000 |
| MOB1 | Cell Signaling | 13730 | 1:1000 |
| pMOB1 (Thr35) | Cell Signaling | 8699 | 1:1000 |
| pYAP (Ser112) | Cell Signaling | 13008 | 1:1000 |
| DUSP6 | Abcam | ab238512 | 1:500 |

**Table 4.** Antibodies used for Immunofluorescence.

| Antibody against | Manufacturer | Cat. No. | Dilution |
| --- | --- | --- | --- |
| cTnT | Thermo Fischer | MA5-12960 | 1:200 |
| cTnI | Abcam | ab47003 | 1:200 |
| Ki67 | Cell Marque | 275R-14 | 1:200 |
| Nestin | Abcam | ab11306 | 1:200 |
| Tomm20 | Cell Signaling | 42406 | 1:200 |
| GJA1 | Abcam | ab11370 | 1:200 |
| αSMA | Sigma-Aldrich | A2547 | 1:400 |
| AurkB | BD Transduction Laboratories | BD 611082 | 1:200 |
| pH3 | Cell Signaling | 9701 | 1:200 |

**Table 5.**
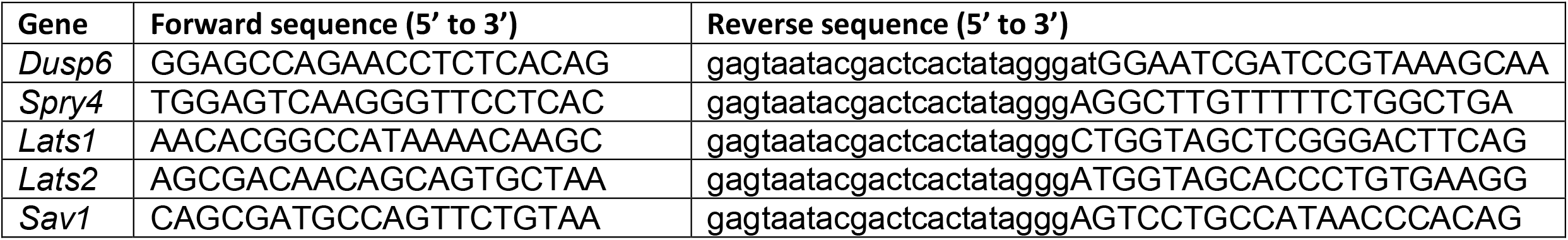
*in situ* probes.

